# Endothelial adaptation to complex flow patterns in a novel in vitro model predicted by computational fluid dynamics

**DOI:** 10.64898/2026.06.27.734995

**Authors:** Stephen B Spurgin, Sepideh Salimi, Vivian S Lee-Kim, Tuli Pramanik, Marcel Mettlen, Hamid Sadat, Ondine Cleaver

**Author notes:** Co-corresponding authors: Ondine Cleaver, Ph.D, Professor, Department of Molecular Biology, University of Texas Southwestern Medical Center, 5323 Harry Hines Blvd., NA8.300, Dallas, Texas 75390-9148, USA, Hamid Sadat, Ph.D., Associate Professor and Associate Chair, Mechanical Engineering Department, University of North Texas, Discovery Park – 3940 N Elm St., F115S, Denton, TX 76207. co-first (Stephen B Spurgin, Sepideh Salimi).

## Abstract

The endothelial cells (ECs) that line blood vessels continuously sense and respond to the physical forces exerted by blood flow. In vivo, pulsatile arterial flow interacts with vessel curvature, branching and other anatomical features to generate complex local hemodynamic environments that dictate the magnitude, direction, pulsatility, and oscillatory nature of wall shear stress experienced by ECs. Currently, accessible and reproducible in vitro models of complex pulsatile flow that recapitulate in vivo vascular anatomy remain limited. Here, we combine a novel rotational-flow endothelial culture platform with detailed computational fluid dynamics (CFD) modeling to characterize four well geometries designed to generate distinct hemodynamic environments. CFD analyses demonstrate that these geometries intrinsically generate pulsatile flow and produce reproducible spatially distinct regions of wall shear stress magnitude, pulsatility, and oscillatory shear within a single culture well. Endothelial alignment mapping and functional assays reveal region-specific cellular responses to the predicted local flow conditions that closely corresponded to the predicted local hemodynamic environment, linking complex flow patterns to endothelial adaptation. The technical advancements of our modeling efforts should support a faster, cheaper, simpler, and—importantly—validated framework for future investigation into EC mechanobiology under complex flow conditions.

**HIGHLIGHTS:**

- Simple engineered well geometries generate distinct hemodynamic microenvironments, mimicking *in vivo* vascular structures, using a conventional orbital shaker.
- Computational fluid dynamics (CFD) reveals spatially distinct patterns of wall shear stress, pulsatility, and oscillatory shear applied to ECs within individual culture wells.
- High average wall shear stress and elevated oscillatory shear index induces a unique perpendicular alignment of ECs to the dominant flow vector.

## INTRODUCTION

The pulsatile flow of blood through branching, curving, and narrowing arteries generates spatially and temporally heterogeneous forces across the vessel wall. Endothelial cells (ECs) lining our blood vessels act as sophisticated mechanotransducers that continuously sense and respond to the variable forces of blood flow.^1,2^ These biomechanical cues are fundamental to both vascular homeostasis and pathological vascular remodeling.^3–8^ Distinct combinations of wall shear stress (WSS), oscillatory shear, pulsatility elicit profoundly different physiological outcomes.^9^ While prior studies have characterized selected flow metrics within orbital shaker systems,^10,11^ a systematic framework linking local multidimensional hemodynamic environments to endothelial cell responses across entire culture surfaces remains lacking.

Steady, unidirectional, laminar flow promotes an anti-inflammatory and atheroprotective endothelial phenotype, whereas disturbed or oscillatory flow triggers pro-inflammatory signaling pathways that predispose vessels to disease.^12^ At the molecular level, these distinct mechanical environments regulate different signaling pathways, such as the protective KLF2/4 axis under stable flow and the pro-inflammatory NFκB pathway under disturbed flow.^13,14^ Disturbed flow also promotes leukocyte recruitment and vascular remodeling during atherogenesis.^15–17^ Finally, EC morphology likewise depends on flow pattern, with oscillatory flow disrupting EC elongation and altering expression of junctional proteins including CDH5 and GJA1.^18^

The complex hemodynamic environments found in blood vessels have been extensively studied using computational fluid dynamics (CFD).^19,20^ Prior CFD work has predicted disturbed and oscillatory flow at arterial branch points and along the lesser curvature of the aortic arch compared to the more laminar flow along the greater curvature or in the descending thoracic aorta.^21,22^ Importantly, these areas of flow disturbance correspond to loss of endothelial alignment to flow as well as acquisition of a pro-inflammatory, atherogenic phenotype.^23^ However, as no two aortas are exactly alike, neither is the pattern of blood flow within them. Even a slight modification of torsion to the aortic arch can result in dramatically different local flow patterns.^24^ Consequently, reproducible *in vitro* systems are needed to precisely relate complex combinations of mechanical input to EC phenotype.

To study this complex mechanobiology in a controlled laboratory setting, the orbital shaker has emerged as a simple, scalable platform for generating fluid motion.^14^ Prior work has utilized CFD to define the shear stress generated in an open well, which yields an area of disorganized flow and low EC alignment in center of the well.^10,25,26^ Additional studies demonstrated that introducing a central post eliminates the area of disturbed flow.^11,27^ One innovative paper generated dramatic pulsatility of flow by tilting the orbital platform.^28^ While CFD analyses have described flow fields in conventional orbital shaker wells, relatively few studies have investigated how geometric modifications such as central posts or added obstacles reshape local hemodynamic environments and endothelial phenotypes.

Despite widespread use of the orbital shaker system, a comprehensive characterization linking local hemodynamic forces to EC responses remains incomplete. Herein, we provide a comprehensive analysis of EC responses to spatially heterogeneous hydrodynamic loads generated by rotational flow culture systems. We performed whole-well immunofluorescence imaging of ECs exposed to flow and related regional cellular responses to the local hydrodynamic environment predicted by CFD simulations. To enable this analysis, we generated detailed maps of WSS, shear stress pulsatility, and oscillatory shear index across the entire culture surface. By linking CFD-derived regional hemodynamic metrics to region-specific endothelial responses, our study establishes a validated framework for interpreting endothelial mechanobiological responses to complex flow environments and provides a foundation for understanding how distinct patterns of disturbed flow regulate endothelial function.

## RESULTS

### Endothelial alignment reflects regional hemodynamics dictated by vessel architecture

The ejection of blood from the heart into an artery (aortic or pulmonary) is a dynamic process. Two key features define the forces exerted by the pulsatile flow of blood on ECs. First, the forces are *time-dependent*, as they change throughout the cardiac cycle, generating varying degrees of pulsatility and/or oscillation of the force (**Fig. 1a**). Second, forces on ECs are *region-dependent*, as the branches, curvatures, and narrowings of blood vessels direct the pulsatile flow of blood to create region-specific flow environments (**Fig. 1b**). These time- and region-dependent forces influence endothelial morphology and function. For example, the lesser curvature of the aortic arch experiences more disturbed and oscillatory flow than the greater curvature of the aortic arch or the descending (straighter) thoracic aorta.^22^ Correspondingly, ECs in the lesser curvature exhibit reduced alignment to flow and a more inflammatory phenotype (**Fig. 1b’**).^23^

**Figure 1.**
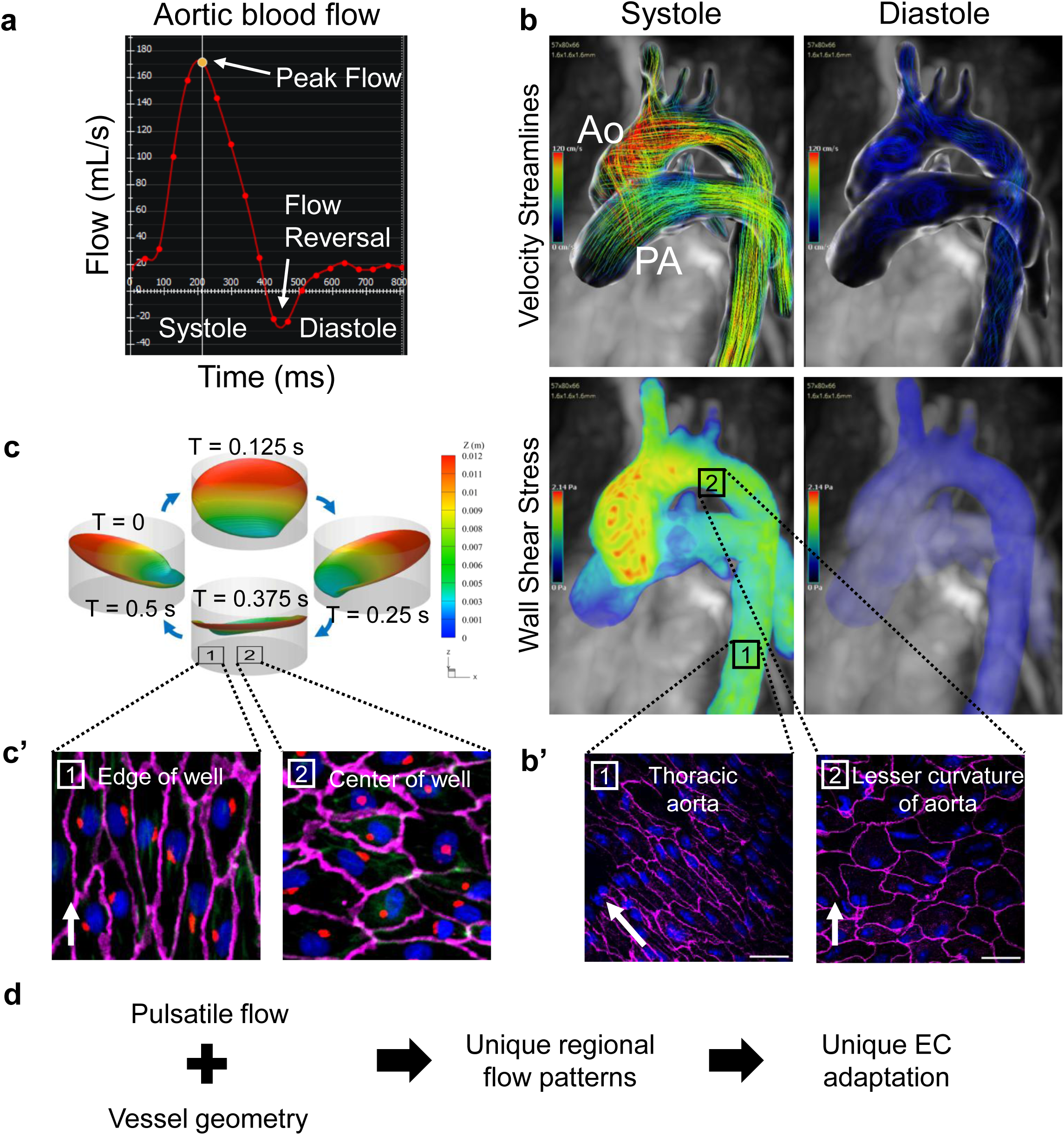
Vessel geometry generates regional hemodynamic environments that impact endothelial alignment, *in vivo* and *in vitro*. Pulsatile arterial blood flow is shaped by the unique curves and branches of blood vessels and generates spatially and temporally unique forces to different regions of the vessel. **a)** Relation of flow velocity to cardiac cycle in a human aorta, illustrating temporal variation in hemodynamic forces experienced by endothelial cells (ECs). **b)** Velocity streamlines from 4D flow cardiac MRI in systole and diastole showing the marked variation in flow pattern throughout the cardiac cycle, with insets **(b’)** showing example of endothelial orientation in selected regions. **c)** Orbital-flow cell culture model showing fluid motion through the rotational cycle of *in vitro* flow, with insets **(c’)** at edge and center of well illustrating EC orientation and alignment in those regions. **d)** Overall schema of EC adaptation to hemodynamic force.

To recreate these hemodynamic conditions *in vitro*, ECs are commonly cultured in standard six-well plates placed on an orbital shaker (**Fig. 1c**). Orbital motion generates fluid flow and shear stress across the endothelial monolayer (**Supplemental Video 1,2,3**). CFD analyses have shown that standard open wells generate predominantly laminar forces near the periphery and disturbed or oscillatory flow in the center.^11^ Consistent with *in vivo* observations, these *in vitro* studies demonstrate that regional flow patterns govern endothelial alignment.^14^ ECs exhibit pronounced elongation and alignment in the outer, laminar flow area, and random cell shape and orientation in the center of the well (**Fig. 1c’**). In sum, these observations demonstrate that the combination of pulsatile flow and vessel geometry creates unique regional flow patterns that influence endothelial cell biology (**Fig. 1d**).

### Design of unique culture well geometries to mimic vascular structures

To reproduce the natural diversity and complexity of *in vivo* vascular geometry, we engineered four distinct configurations for a well of a six-well plate (**Fig. 2a**). Two previously described models served^10,25,28^ as reference models: an unmodified well (the “open” well, Case 1), and a central post well that forms an annular, fluid-filled channel (the “donut” well, Case 2). Building upon these established designs, we developed two additional geometries. In Case 3, the central post was displaced laterally to create an asymmetric annular channel. We hypothesized that this geometry would create two features: A) a region of flow acceleration through the narrow side of the post, and B) a region of flow oscillation downstream of the narrow channel (the “offset” well, Case 3). Finally, we modified Case 2 with a curved insert to create flow compression upstream and flow disturbance downstream of the obstacle (the “obstacle” well, Case 4).

**Figure 2:**
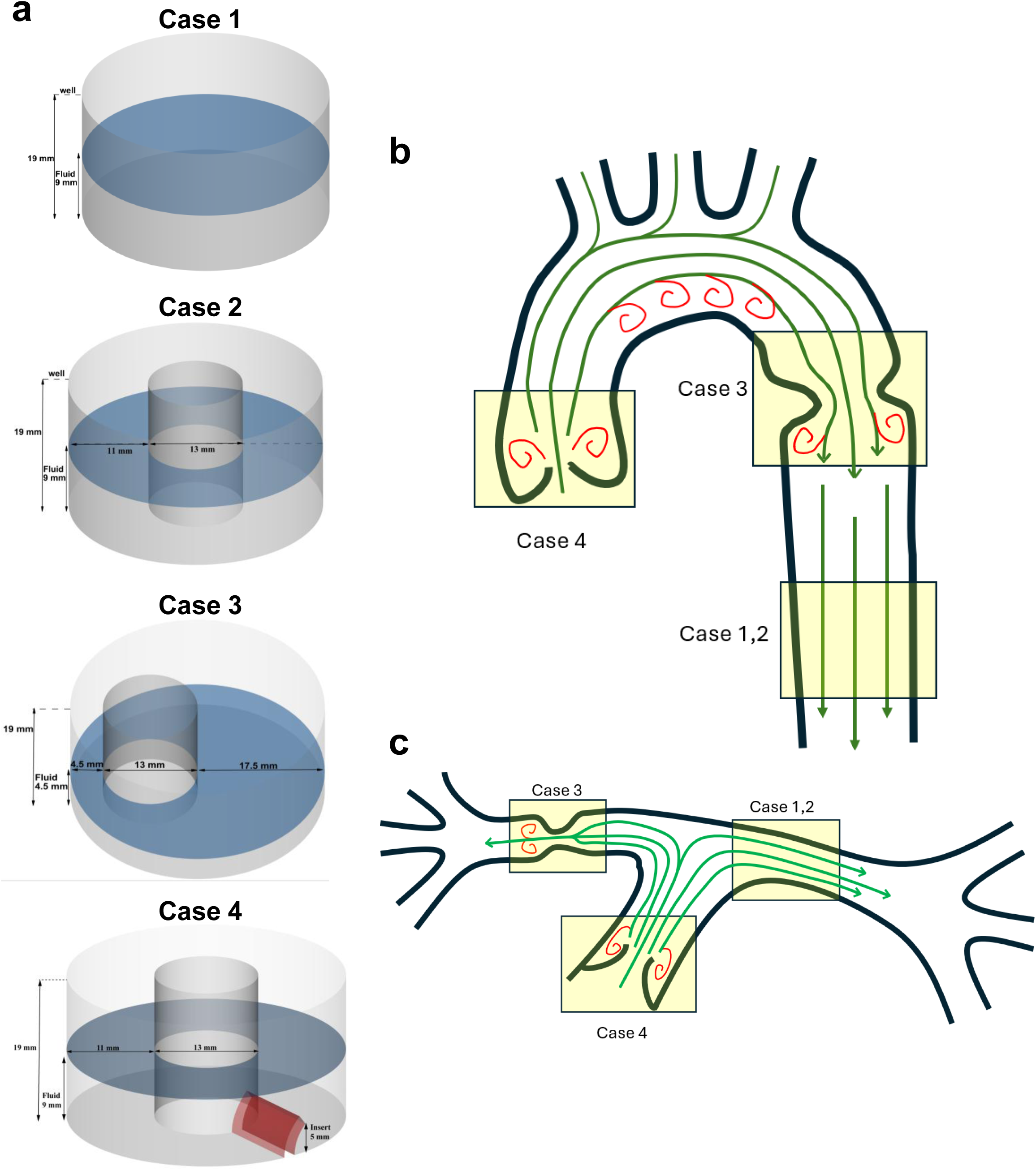
Modeling complex physiologic and pathologic vascular flow environments *in vitro*. **a)** Representative 3D models used for CFD modeling of culture well geometries used to generate distinct flow environments: open (empty), donut (central post), offset (displaced post), and convex obstacle. **b)** Schematic of aortic blood flow. General path of blood flow shown in green, with previously identified regions of turbulent or oscillatory flow noted in red. Aortic coarctation is is illustrated as a representative stenotic lesion corresponding to the Case 3 geometry. **c)** Schematic of the pulmonary arterial blood flow. Regions with disturbed flow are highlighted in red. A stenotic right pulmonary artery (RPA) is shown as an example of a flow-limiting lesion represented by the Case 3 geometry.

To facilitate direct comparison among the different geometries, the fluid depth was standardized in Case 1, Case 2, and Case 4 to maintain continuous flow over the obstacle in Case 4. In Case 3, the fluid depth was reduced by 50% to prevent overflow from the top of the well adjacent to the narrow constriction during orbital motion.

These geometries were designed to reproduce hemodynamic conditions encountered in several clinically relevant vascular settings (**Fig. 2b,c**). The more cohesive, laminar flow of Case 2 would mimic the flow observed in large conduit vessels such as the descending aorta or branch pulmonary artery after making the turn toward the lung. Case 3 mimics flow through a vascular stenosis, generating both accelerated flow within the constriction and disturbed post-stenotic flow, similar to that observed in aortic coarctation or branch pulmonary artery stenosis. Finally, the flow across the stiff, convex curvature of the obstacle in Case 4 models the disturbed flow across a stiff/thickened semilunar (aortic or pulmonary) valve leaflet. For each case, we generated detailed models for analysis by CFD (**Supplemental Fig. 1**).

### Impact of fluid depth and velocity on local wall shear stress

Because viscous fluids satisfy the “no-slip” condition, fluid immediately adjacent to the moving well surface travels with the wall. Thus, the fluid velocity relative to the wall is zero at the wall itself. Consequently, velocity values directly on the well surface would be uniformly zero and would not reveal the underlying flow field experienced by the ECs. Therefore, fluid velocity is reported on a plane located slightly above the well surface, whereas WSS, which depends on the velocity gradient normal to the wall, is measured directly on the well surface.

During orbital rotation, fluid depth varied dynamically, changing nearly four-fold between the outer (high) and inner (low) regions of the orbit (**Fig. 3a**). Velocity contours evaluated at the liquid–air interface and on a plane located 2 mm above the bottom surface revealed complex spatial flow patterns that evolved throughout the orbital cycle **(Fig. 3b,c)**. Greater fluid depths near the outer perimeter of the well promoted the formation of vortical flow structures, whereas shallower regions exhibited more organized flow.

**Figure 3:**
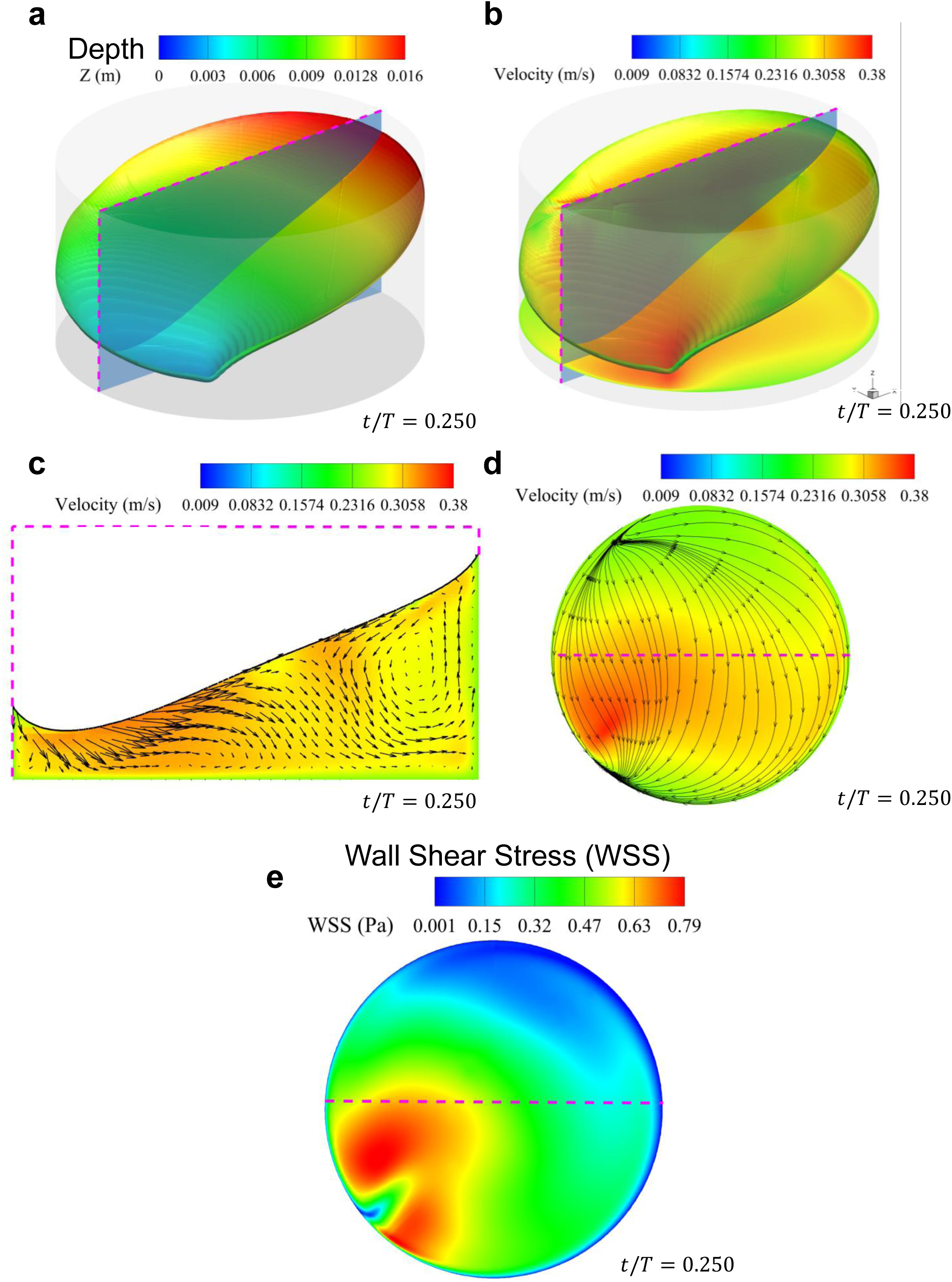
Fluid depth and velocity at a representative point of the orbital cycle in the open well. At a single time point, CFD was used to calculate multiple parameters, including **(a)** fluid depth, **(b)** fluid velocity magnitude at the fluid-air interface and 2mm above the bottom of the well, **(c)** velocity vector profile in a cross-section extending from the shallowest to the deepest region of the well, **(d)** velocity vector profile directly above the bottom of the well, and **(e)** WSS distribution across the culture surface.

These variations in fluid depth strongly influenced the velocity gradient at the wall and, consequently, the local WSS. Although regions of similar near-wall flow velocity were observed throughout the well, shallower fluid layers generated steeper velocity gradients and therefore higher WSS than deeper regions. As a result, elevated WSS did not necessarily coincide with the highest local flow velocity. Instead, the combined effects of local fluid depth and flow velocity produced the spatial distributions of near-wall velocity **(Fig. 3d)** and WSS **(Fig. 3e)** observed throughout the orbital cycle.

The offset well further illustrates these principles. As the deeper fluid layer accelerated through the narrow constriction, elevated WSS developed despite the increased local fluid depth **(Supplemental Fig. 2a–c)**. Notably, elevated WSS arose through two distinct mechanisms: (1) acceleration of flow through the constriction (pink arrow) and (2) shallow fluid depth despite lower instantaneous flow velocity (yellow arrows). The accelerated flow within the constriction was accompanied by increased vorticity **(Supplemental Fig. 2d)**, whereas a separate band of elevated WSS developed along the opposite edge of the well, where relatively organized laminar flow occurred under shallow fluid conditions **(Supplemental Fig. 2d’, c)**.

### Distinct well geometries generate unique patterns of WSS

Incorporating local depth and velocity relative to the motion of the well, CFD simulations were used to calculate the WSS across the surface of the well. For each case, we generated heatmaps of the average WSS (**Fig. 4a**), as well as recordings of the instantaneous WSS throughout the cycle (**Supplementary Videos 4-7**). To illustrate the directionality of the net applied shear, the net WSS vector within the plane of the culture surface is overlaid on each heatmap, with vector length proportional to the magnitude of the net shear stress. The distinction between the time-average WSS magnitude and the magnitude of the net WSS vector is shown in **Supplementary Fig. 3.** Additionally, the wall shear stress gradient (WSSG) was calculated for each case (**Supplementary Fig. 4**).

**Figure 4:**
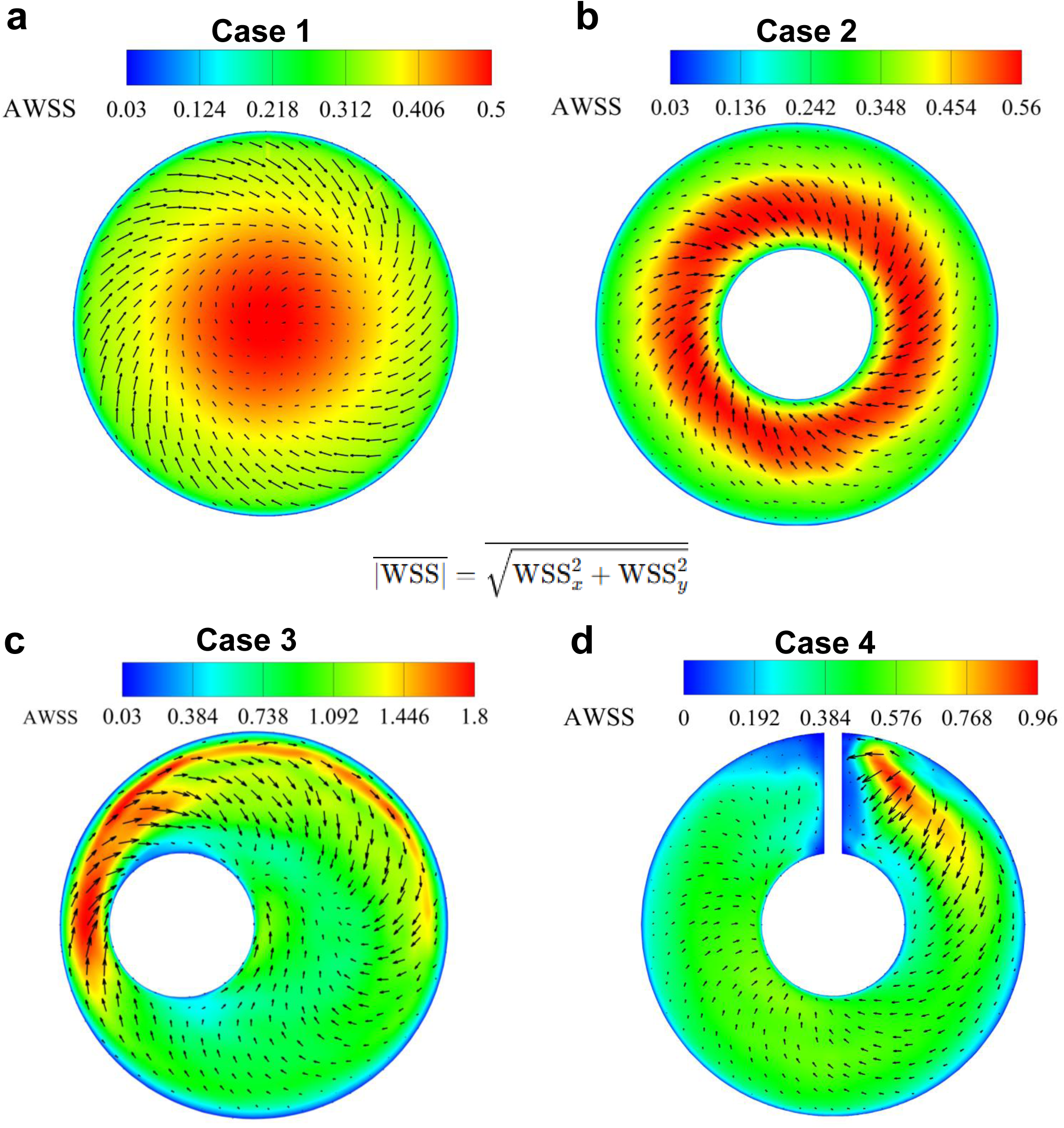
Distinct rotational flow geometries generate unique spatial patterns of wall shear stress (WSS). For each case **(a-d)**, heatmaps show the time-averaged magnitude of the WSS across the culture surface, displaying the average intensity of shear stress (AWSS) experienced each location throughout the orbital cycle. Overlaid arrows indicate the direction of the net WSS vector, with arrow length scaled to its magnitude. Heatmaps and vector fields illustrate both the spatial distribution and predominant direction of shear stress within each well geometry.

The four well geometries generated distinct spatial distributions of both WSS magnitude and net shear direction. For all cases, net shear vectors typically angled inward with respect to the direction of fluid flow. As expected, the open well case (Case 1) exhibited stronger net vectors at the outer edge of the dish, with weaker net forces toward the center. However, it is important to note that while the *net* shear vector in the center of the well is low, the average WSS *magnitude* is highest in that region. The annular geometry of donut well (Case 2) eliminated this central region of undirected, high magnitude, and low net shear, resulting in the most uniform distribution of shear vectors among all four geometries. The highest average WSS values of any case were found in the narrow gap of the offset configuration (case 3), which generated an average WSS of 18 dyn/cm^2^. This contrasted with the opposing areas of the same well where average WSS values were typically around 8 dyn/cm^2^. Finally, the obstacle well (Case 4) yielded high average WSS values downstream of the obstacle but surprisingly uniform average WSS throughout the rest of the well.

### Pulsatile flow generated by orbital shaker in cell culture

Given the dynamic changes in fluid depth and velocity that occur throughout the orbital cycle, we next characterized the temporal variability of WSS within each well geometry. WSS throughout the cycle is shown in **Supplemental Video 2**. To illustrate the temporal characteristics of the local shear environment, WSS waveforms were generated at representative locations (**Fig. 5a**) by plotting the magnitude of WSS as a function of time over a complete orbital cycle (**Fig. 5b**). Distinct temporal fluctuations in WSS were observed at most locations, in all configurations, demonstrating that ECs are exposed to spatially heterogeneous pulsatile shear environments.

**Figure 5:**
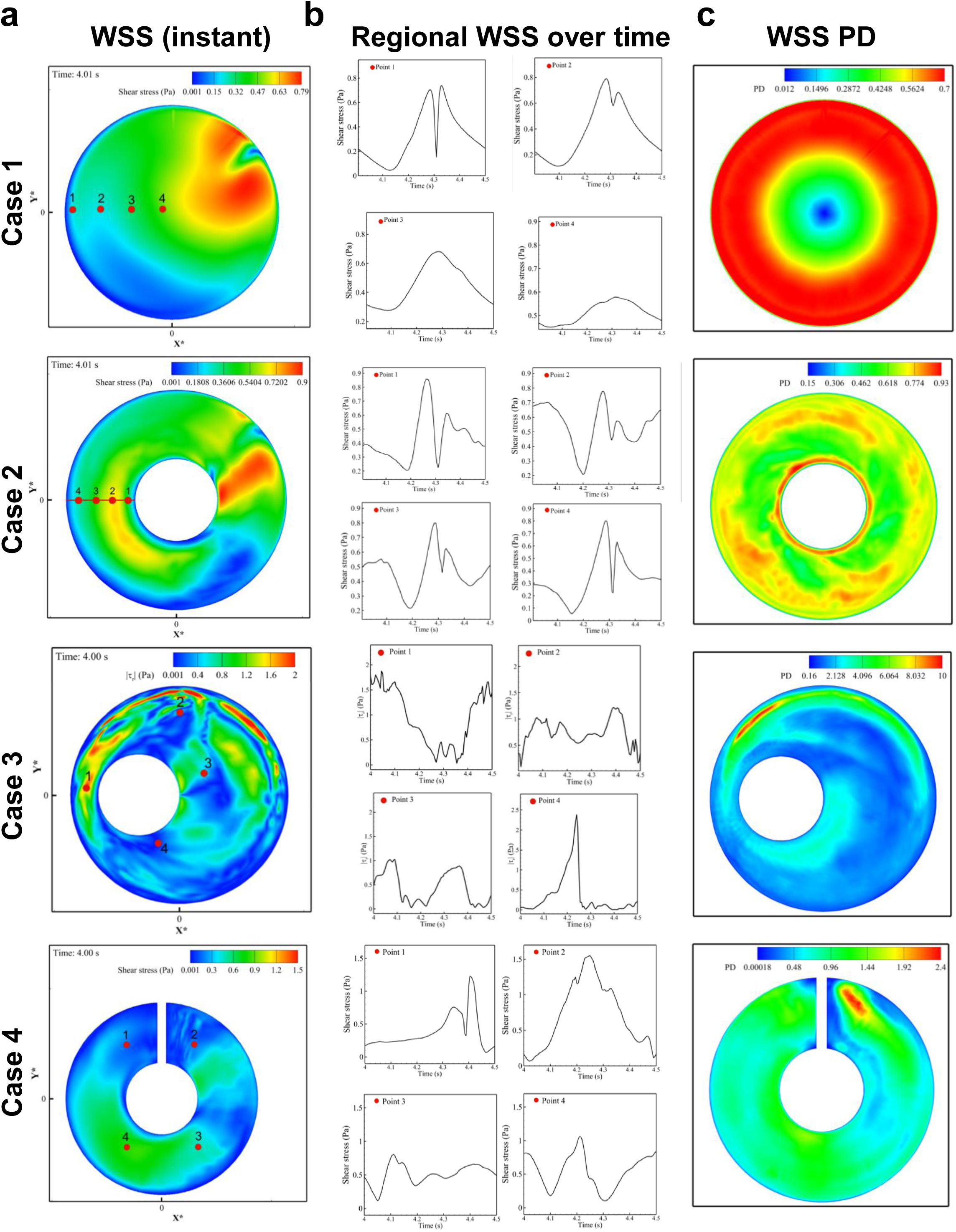
Shear stress magnitude is pulsatile within rotational flow models. **a)** WSS magnitude at t/T = 0 for Case 1-4 well configurations. Four points are shown with corresponding graphs **(b)** of the shear stress throughout the orbital cycle. **c)** WSS-driven Pulse Difference (PD) for each well geometry, showing spatial distribution of pulsatility.

The temporal characteristics of these waveforms varied substantially among the four geometries. In the open well, we found pulsatility was prominent in the outer portion of the well, but reduced to almost zero pulsatility at the center (Case 1, point 4). The donut well exhibited a highly stereotyped pulsatile waveform, characterized by a dip, followed by two distinct peaks before returning to baseline. Among all geometries, the offset well displayed the greatest overall pulsatility, compared to the other three cases, with peak values occurring downstream of the “stenosis.” This case also contained the sharpest spatial transition in pulsatility, with a relatively sheltered area upstream of the constriction (Case 3, point 4). The obstacle well likewise generated pronounced pulsatility, particularly downstream of the obstacle (Case 4, point 2).

Next, to quantify spatial variations in pulsatility throughout the well, we generated heatmaps of WSS pulse difference (WSS PD) for each well. WSS PD (WSS PD = WSSmax – WSSmin) was approximately 7 dyn/cm^2^ throughout much of the open and donut wells (**Fig. 5c**). The offset well (Case 3) exhibited the greatest pulsatility, with average WSS PD values in the 20 to 40 dyn/cm^2^ range and reaching a peak PD approaching 100 dyn/cm^2^. The obstacle well (Case 4) also demonstrated elevated pulsatility throughout, with a peak pulsatility downstream of the obstacle.

To facilitate comparison with commonly used clinical metrics,^29^ we also calculated the WSS Pulsatility Index (PI) (PI = [WSSmax – WSSmin]/WSSmean) for each case (**Supplemental Fig. 5a**). PI revealed patterns like those observed with PD, but further emphasized area 4 in Case 3, where low mean WSS amplified the relative pulsatility. Finally, we calculated the root means squared (RMS) WSS, a measure of the magnitude of WSS deviations from the local mean value over the orbital cycle (**Supplemental Fig. 5b**). RMS WSS is therefore another measure of shear pulsatility. Collectively, these analyses demonstrate that different well geometries generate spatially distinct pulsatile fluid microenvironments, exposing neighboring endothelial populations to markedly different temporal patterns of shear stress.

### Quantification of oscillatory shear stress in two dimensions

To quantify the impact of oscillatory shear stress across each well geometry, we calculated the oscillatory shear stress index (OSI) for the surface of each well. Under the standard definition (**Supplementary Table 1**), OSI ranges from 0 to 0.5, where 0 reflects fully unidirectional shear and values approaching 0.5 indicate increasingly bidirectional or multidirectional shear with minimal net directional bias over the cycle.

In a blood vessel, the primary axes of WSS can be described relative to the longitudinal and circumferential axes of the vessel wall (**Supplementary Fig. 6a, a’**). Analogously, in our rotational cell culture model, the direction of WSS can be resolved into radial (r) and tangential (θ) components, corresponding to side-to-side and back-and-forth motion relative to the overall direction of flow (**Supplementary Fig. 6b-d**). Although OSI is conventionally reported as a single scalar quantity, decomposing OSI into its radial and tangential components provides additional insight into the multidirectional nature of the local flow environment.

The relative contributions of radial and tangential oscillation differed markedly among the four geometries (**Figure 6**). The open well (Case 1) exhibited very high composite OSIrθ throughout the culture surface, with higher values in the center of the well (**Fig. 6a**). In the central region, there is little directional predominance to the OSI, as neither tangential nor radial components dominated (neither OSIr nor OSIθ is zero), resulting in a highly multidirectional flow environment. This represents flow that is predominantly swirling. In the donut well (Case 2), oscillatory shear was present throughout the culture surface, which is primarily due to tangential rather than radial oscillations (**Fig. 6b**). The offset well has very low OSI in the narrow gap but marked OSI in an area downstream and behind the post with respect to the direction of flow (**Fig. 6c**). Finally, the obstacle well (Case 4) displayed a remarkable spatial asymmetry in OSI (**Fig. 6d**)—high OSI upstream of the obstacle, and low OSI downstream, once past an initial area of flow recirculation (**Supplemental Fig. 7**). These analyses demonstrate that similar OSI values can arise from fundamentally different combinations of radial and tangential oscillation, revealing distinct multidirectional flow environments generated by different well geometries.

**Figure 6:**
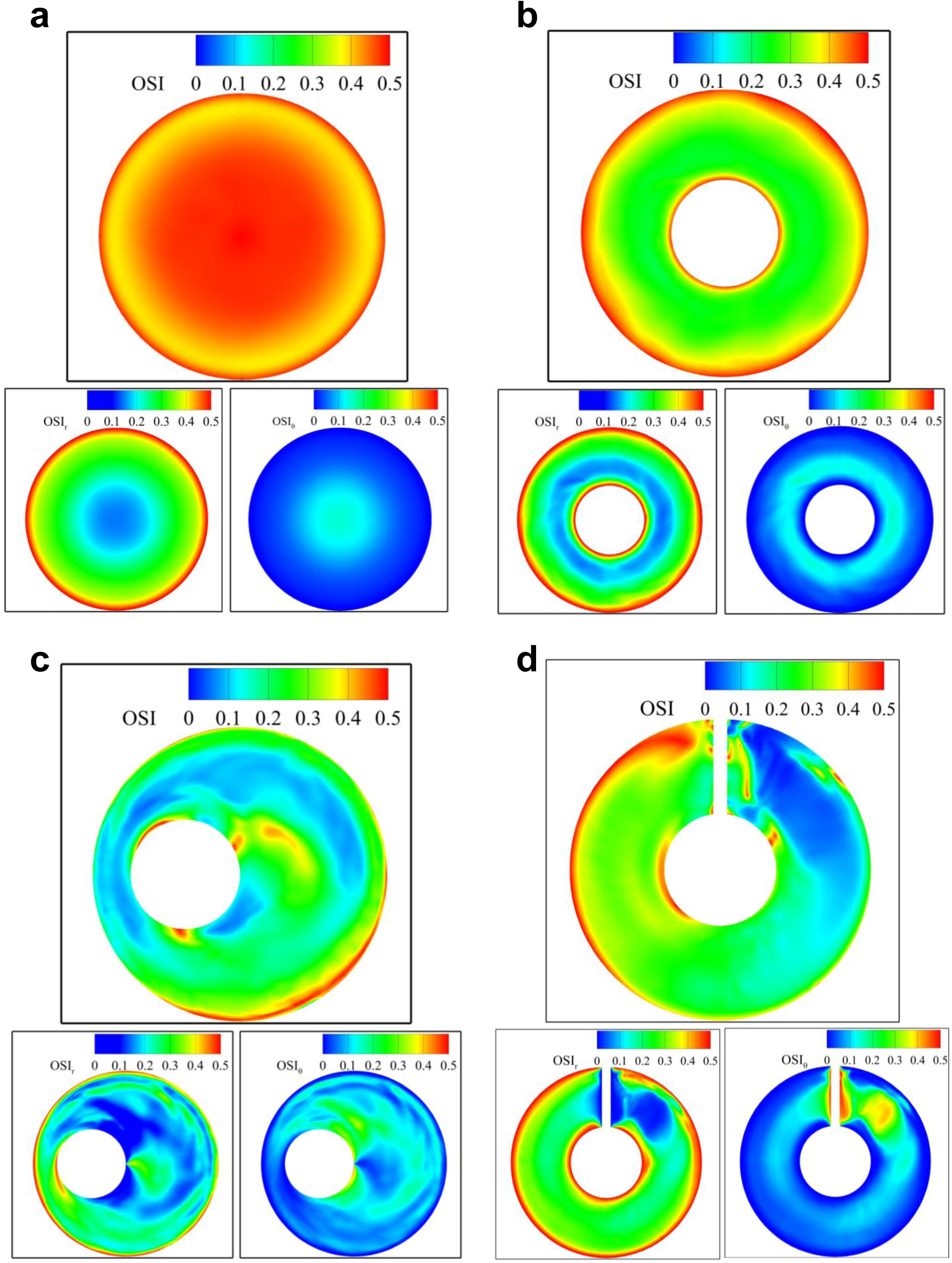
Two-dimensional decomposition of oscillatory shear stress index (OSI) in rotational flow. OSI was calculated by decomposing oscillatory shear into radial (r) and tangential (*θ*) components, which are displayed separately below the composite OSI maps. Results are shown for open well **(a)**, donut **(b)**, offset **(c)**, and obstacle **(d)** well geometries. Composite OSI maps illustrate overall oscillatory behavior, whereas the radial and tangential maps identify the dominant directional components contributing to the local oscillatory shear. Values range from 0 (fully unidirectional shear) to 0.5 (maximally oscillatory shear). Here, *OSI* = *OSI*_rθ_.

### Local cellular alignment is dictated by net WSS vector throughout the well

Whole-well confocal imaging revealed a close relationship between local hemodynamic conditions and endothelial organization. Cell borders, Golgi, nuclei, and actin cytoskeleton were visualized by immunofluorescence (IF) using CDH5, GM130, DAPI, and phalloidin, respectively, and both regional and cell-specific morphology was analyzed (**Supplemental Fig 8**). Whole well imaging at 10x magnification enabled direct comparison of regional endothelial organization with the corresponding CFD-derived maps of the time-averaged WSS vector **(Supplemental Fig. 9a)** and oscillatory shear index (OSI; **Supplemental Fig. 9b**).

In the open well (Case 1), ECs aligned closely with the net WSS vector in the outer portion of the culture surface. There is a point of convergence of the shear vectors (**Supplemental Fig 9. Case 1, white dotted circle**), where the net shear vector transitions from “angled inward” to “angled outward” with respect to the clockwise motion of the fluid in the well. This also corresponds to a point of transition from an OSIrθ of 0.4 toward 0.5. Notably, ECs remained highly elongated in regions of high average WSS and high OSIrθ, yet no longer aligned with the net WSS vector. Complete loss of EC elongation and alignment was observed only in the small area at the very center where no dominant net shear vector existed (**Supplemental Fig. 10**).

Similar alignment patterns were observed in the donut well (Case 2), where ECs aligned with the inwardly angled net WSS vector throughout most of the annular channel. Alignment was lost only at the outer boundary, consistent with regions of elevated OSI and low net shear. In the offset well (Case 3), ECs aligned strongly in both high- and low-WSS areas, but lost alignment at two locations: the region of high OSI, where the two WSS vector fields collide downstream of the “stenosis”, and upstream of the “stenosis” where WSS vectors diverged adjacent to the offset post at the entrance to the narrow gap. Finally, in the obstacle well (Case 4), there was strong alignment to flow downstream of the obstacle in regions of low OSI, whereas alignment was disrupted upstream of the obstacle, where oscillatory shear was greatest. Overall, the cellular alignment throughout the well largely matches the average WSS vector, except in areas of high oscillations of WSS.

To investigate EC responses to the combination of high OSI and high WSS, we returned to the classic open-well configuration. Four key areas were selected for analysis (**Fig. 7a-A’’**). We found good alignment between the cell major axis and the WSS vector (**Fig. 7b,c**) in the outer two selected areas: areas with lower OSI and low to medium average shear stress. By contrast, area 3 (high WSS, high OSI) revealed a striking perpendicular alignment relative to the net WSS vector. Area 4—the exact center of the well—demonstrated loss of cellular alignment, in keeping with the loss of a net WSS vector. The Nuclear-to-Golgi (NtG) axis was oriented approximately -30°/+150° from the net WSS vector in areas 1 and 2, while there was no clear directionality of the NtG in areas 3 and 4, where ECs experience higher OSI values (**Fig. 7d**).

**Figure 7:**
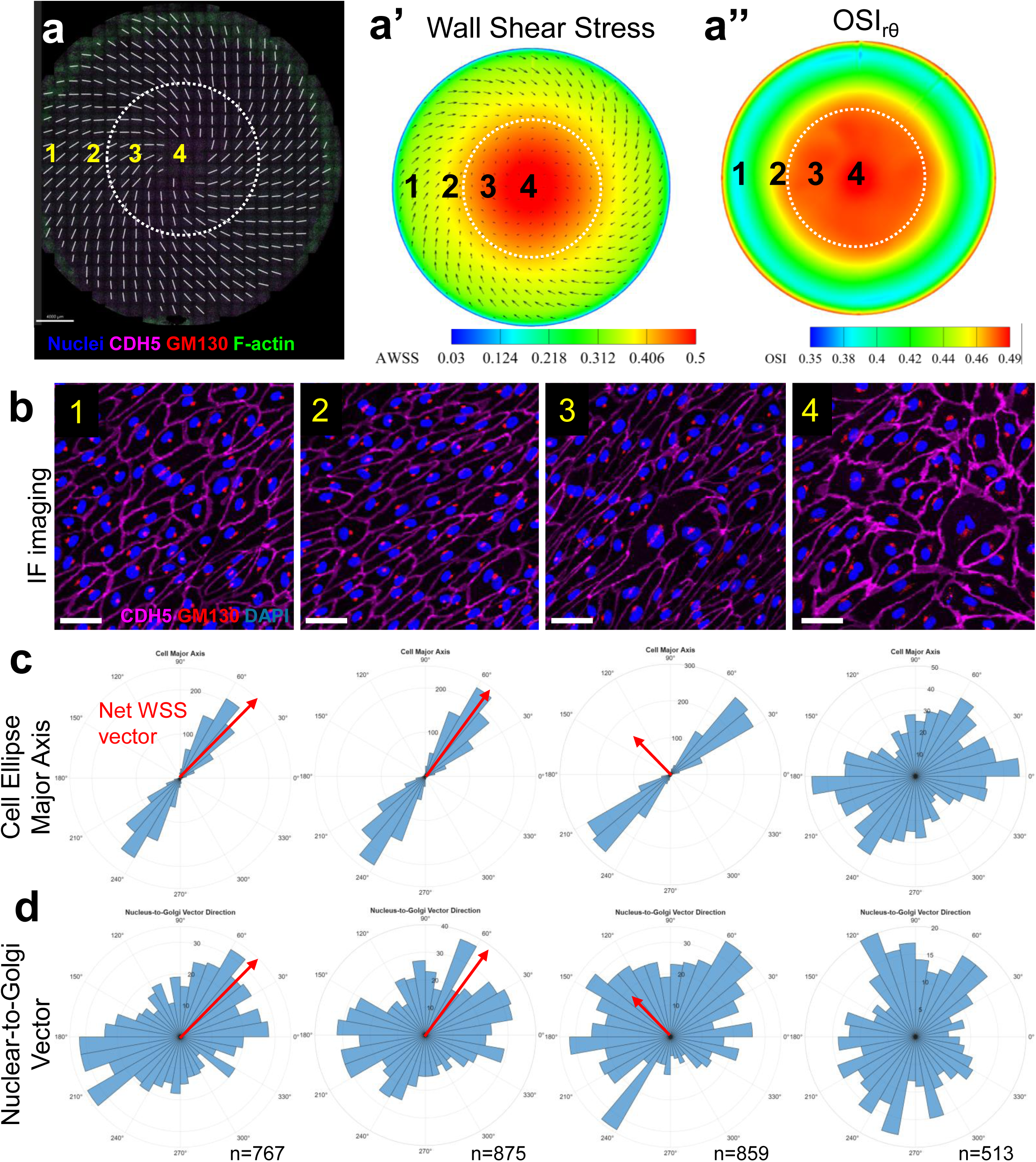
Regional endothelial alignment transitions in regions of elevated oscillatory shear in the open well. Whole-well immunofluorescent imaging for markers as indicated **(a)**, average WSS magnitude heatmap with net WSS vector overlay **(a’)**, and composite OSI_rθ_ map **(a’’)** for the open well. The dotted white line denotes the transition point where AWSS, net WSS vectors, and OSI value. **b)** Key regions are numbered 1-4, with representative close-up images of IF stains for CDH5 (cell borders), GM130 (Golgi apparatus), and DAPI (nuclei). **c)** For each region, over 500 cells were analyzed, and the orientation of the cell major axis is shown in polar histograms. Note that cell major axis is directionless and so the 180° mirror image is also graphed. The net WSS vector is indicated by the red arrow. **d)** For the same regions, a vector from the center of the nucleus to the center of the Golgi are shown. Regions 1 and 2 exhibit alignment parallel to the net WSS vector, whereas Region 3 demonstrates perpendicular alignment under conditions of elevated WSS and OSI.

To further investigate the relationship between cellular alignment and local applied shear forces, we selected key locations in the offset well for analysis (**Fig. 8a-a’’**). In regions of low OSI and moderate-to-high WSS the major axis of cells aligned closely with the average WSS vector (**Fig. 8b,c**). Unexpectedly, in our assay (HPAECs, 5 days of flow) the NtG axis was oriented approximately +/- 90° relative to both the cell major axis and net WSS vector (**Fig. 8d**). This contrasts with prior studies describing Golgi polarization to be either upstream relative to flow, or downstream relative to migration.^30^ Similar to the high WSS/high OSI regions of the open well, region 3 of the offset well demonstrated perpendicular alignment to the mean WSS vector. Interestingly, the NtG axis in region 3 was largely in-line with the mean WSS vector, indicating a position of Golgi downstream of flow. Region 4 of the offset well (medium WSS and low OSI) demonstrated alignment of both cell major axis and perpendicular alignment of NtG, as seen in region 1.

**Figure 8:**
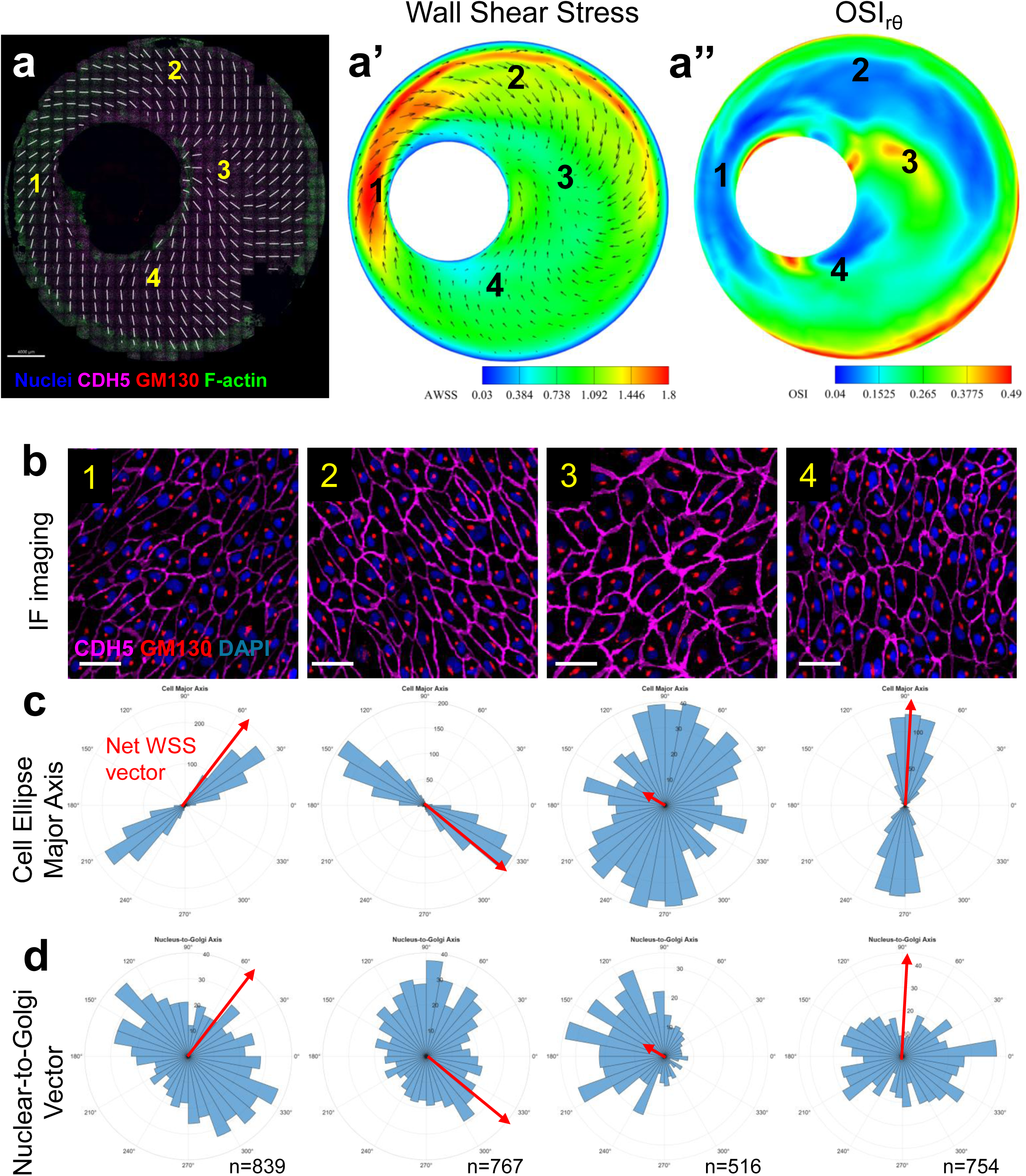
Regional endothelial alignment follows local hemodynamic direction in the offset well: Whole-well immunofluorescent imaging for markers as indicated **(a)**, average WSS (AWSS) magnitude heatmap with net WSS vector overlay **(a’)**, and composite OSI_rθ_ map **(a’’)** for the offset well. **b)** Key regions are numbered 1-4, with representative close-up images of IF stains for CDH5 (cell borders), GM130 (Golgi apparatus), and DAPI (nuclei). **c)** For each region, more than 500 cells were analyzed, and cell major-axis orientation is shown in polar histograms. Note that cell major axis is directionless and so the 180° mirror image is also graphed. The net WSS vector is indicated by the red arrow. **d)** For the same region, vectors from the center of the nucleus to the center of the Golgi are shown.

### Distinct cytoskeletal adaptations of endothelial cells to unique patterns of force

We next investigated how distinct hemodynamic forces influence endothelial cell shape/alignment and cytoskeletal adaptations. The offset well was selected for detailed analysis because it yields critical and variable areas of high/low shear, pulsatility, and OSI. Basic quantifications of nucleus circularity, cell area, and the length of the cell major axis revealed larger, unaligned cells with rounder nuclei in region 3 relative to regions 1, 2, and 4, in keeping with region 3’s high OSI (**Supplemental Fig. 11**). We note increased actin intensity under high shear/high pulsatility area of region 1, with formation of prominent stress fibers (**Fig. 9a,b**). In regions of moderate WSS and low OSI (regions 2 and 4), the actin closely approximates the cell borders. However, in region 3, which experiences moderate WSS and high OSI, we observe lack of cell elongation, wider bands of actin at the cell borders, and occasional actin punctae that are also localized near the EC border (arrows). Quantification of average actin intensity showed significant differences by region, with highest expression in the high shear region 1. CDH5 intensity, a crucial cell-cell junction protein specific to endothelial cells, was highest in the disorganized ECs in region 3 of Case 3 (**Fig. 9c**).

**Figure 9:**
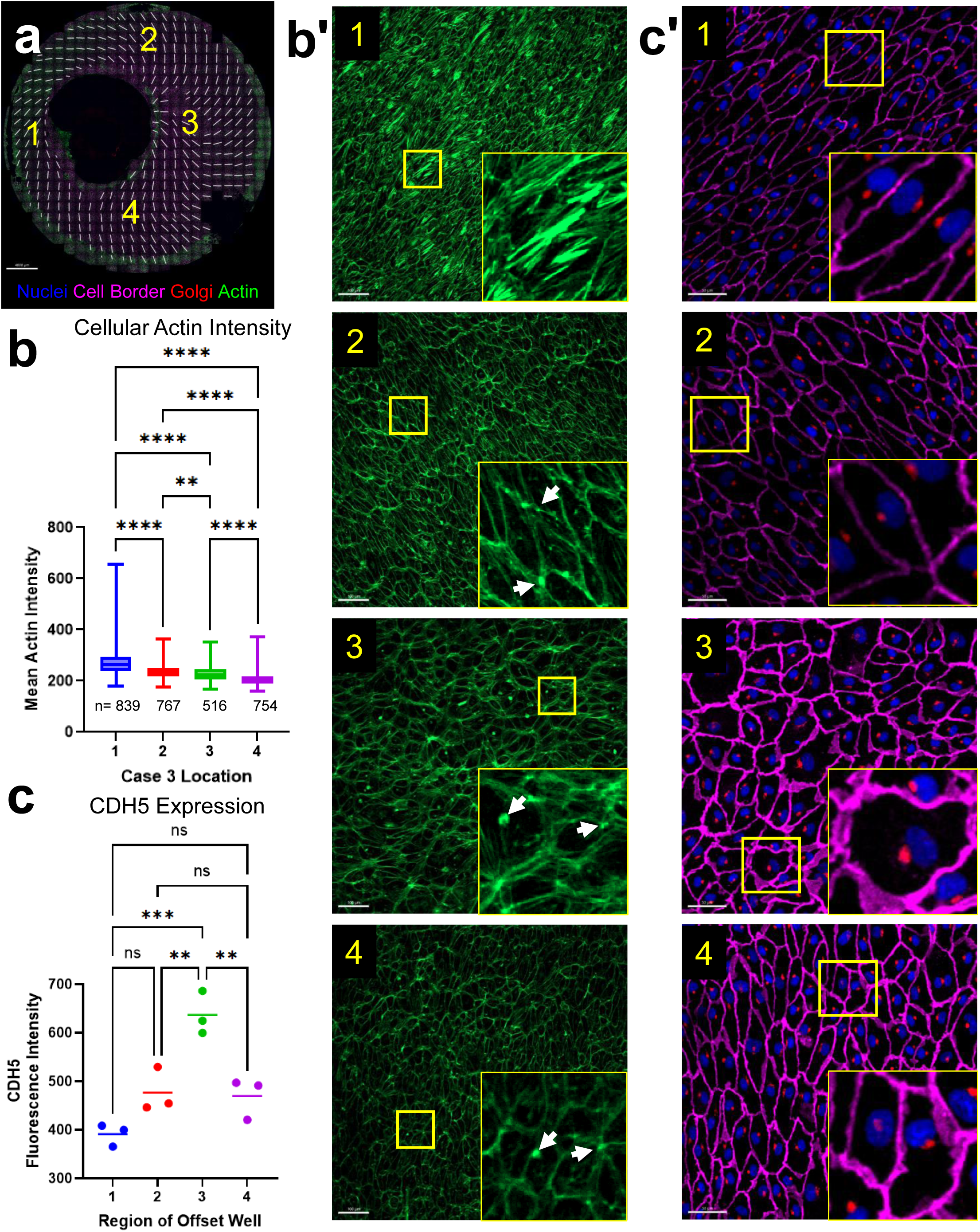
Characterization of actin morphology in the offset well. **a)** Regions selected for analysis. **b,b’)** Quantification of cellular actin expression by phalloidin staining, with insets shown for specific regions analyzed. Number of cells analyzed is listed in the graph. Scale bar is 100 µm. **(c,c’)** Quantification of cell-cell junctions by CDH5 intensity, in the indicated regions of three different Case 3 wells. Scale bar is 50 µm. Groups in (b) and (c) were compared by one-way ANOVA.

Analysis of intensity and pattern of F-actin organization within the ECs in different regions of the obstacle well (**Fig. 10a,b**) further revealed three broad types of cytoskeletal phenotypes associated with different combinations of WSS and OSI. In areas of high shear and low OSI, we observed high actin expression and prominent stress fibers aligned with the direction of flow (**Fig. 10, region 1**). Under conditions of low shear and low OSI, ECs still aligned to the shear and displayed moderate F-actin expression but lacked well-defined stress fibers (**Fig. 10, region 2**). By contrast, areas experiencing high OSI and low WSS displayed increased F-actin expression without organization into shear-aligned stress fibers (**Fig. 10, region 3**). Together, these data suggest that WSS magnitude and oscillatory shear contribute differently to cytoskeletal remodeling, with high WSS promoting stress-fiber formation and high OSI disrupting their directional organization.

**Figure 10:**
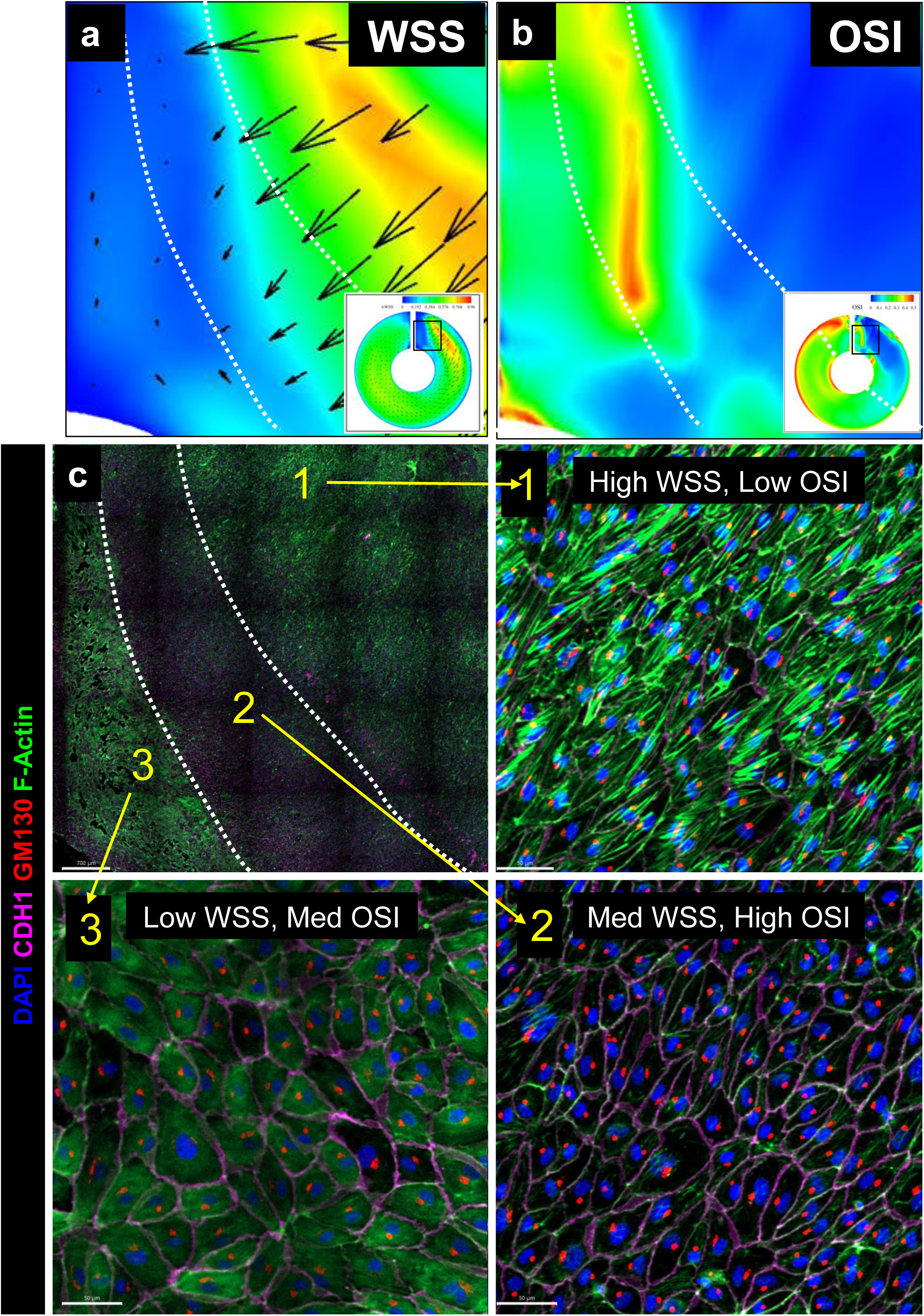
Distinct cytoskeletal phenotypes arise in response to variable local WSS and OSI conditions in a single well. The area downstream of the obstacle in Case 4 exhibits a gradient of WSS **(a)** and OSI **(b)** values, creating distinct local hemodynamic environments for comparison. **(c)** Representative F-actin organization (stain with phalloidin) in regions exposed to distinct combinations of WSS and OSI. High WSS and low OSI (area 1) promote formation of prominent shear-aligned actin stress fibers. Regions with high OSI (area 2) exhibit little EC alignment and no stress fibers. Regions with low WSS and medium OSI (area 3) show higher F-actin expression but no EC alignment.

### Monocyte adhesion in regions of disturbed flow reveal localized EC activation

The lesser curvature of the aortic arch experiences higher OSI, attracts resident macrophages, and is a known site of EC activation and subsequent atherosclerotic plaque formation in mice (**Fig. 11a**).^23^ To further validate the biological relevance of our rotational flow cell culture model, we asked whether ECs exposed to high OSI exhibited an inflammatory EC phenotype that promote monocyte adhesion. Monocyte adhesion was consistently increased in regions of high OSI in all wells (**Fig. 11b,c**). In the open well, for instance, area 2 displayed over 10-fold higher macrophage adhesion than area 1. Together, these findings indicate that regions of disturbed flow within the culture wells promote EC activation and acquisition of a pro-adhesive phenotype.

**Figure 11:**
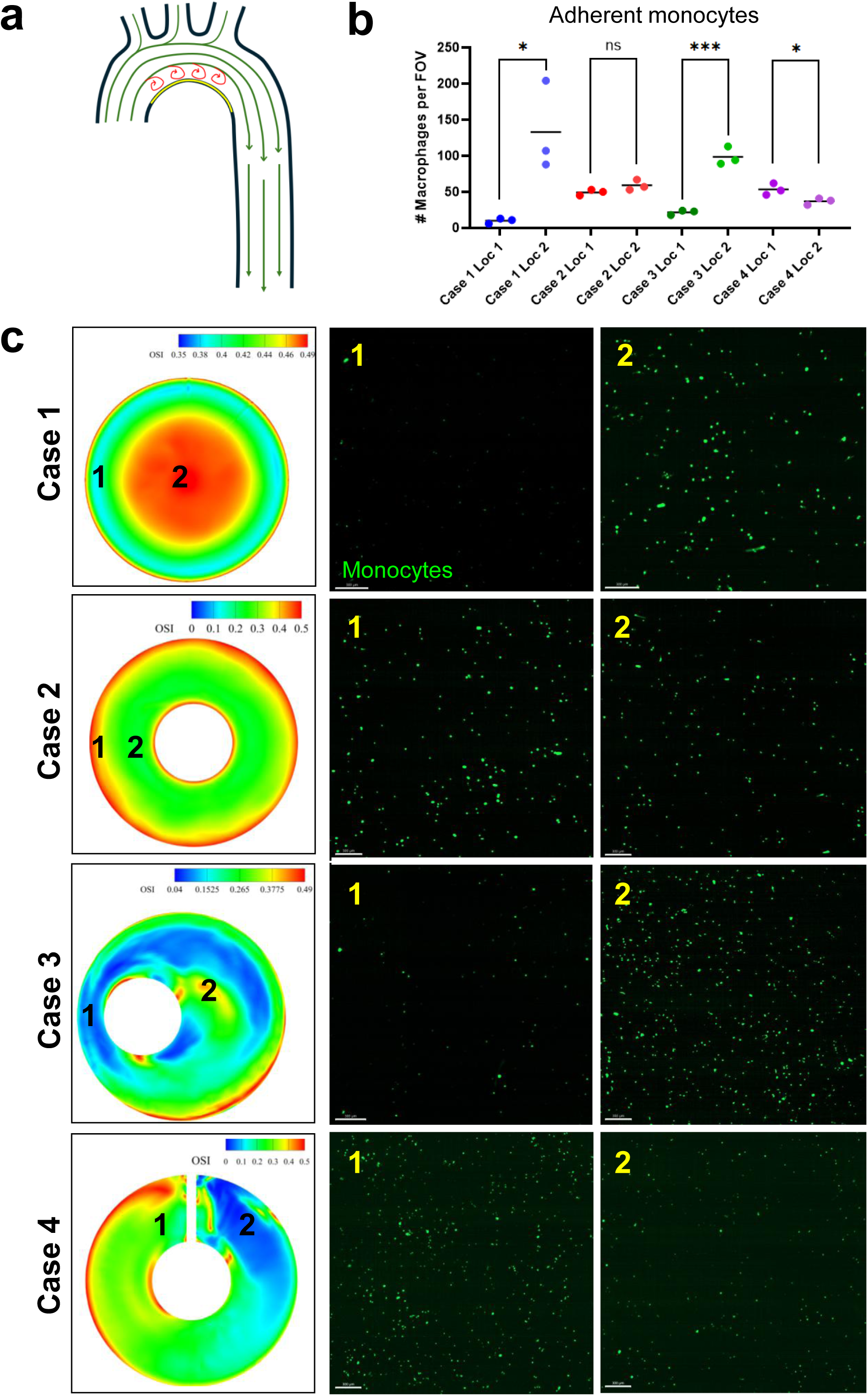
Increased monocyte adhesion to HAECs in regions of oscillatory flow. **a)** The lesser curvature of the aortic arch exhibits elevated oscillatory shear and is a well- established monocyte recruitment, endothelial activation, and atherosclerotic plaque formation *in vivo*. **b)** Quantification of monocyte adhesion in regions (in c) *in vitro* that exhibit low and high regions of oscillatory WSS. **(c)** Representative regions and fluorescence images showing adherent fluorescently-labeled monocytes (green Calcein AM) atop endothelial monolayer. Regions exposed to elevated oscillatory shear exhibited increased monocyte adhesion, consistent with localized endothelial activation under disturbed flow conditions. Student’s t-test was used to compare groups.

## DISCUSSION

The crucial role of hemodynamics in conditioning endothelial cell biology requires highly defined *in vitro* systems to accurately investigate EC responses. Here, we combined CFD and endothelial morphology analysis to characterize the hemodynamic forces in the rotational flow model of cell culture. By integrating engineered well geometries, CFD, and whole well endothelial phenotyping, we transform a conventional orbital shaker system into a practical platform for generating and validating multiple physiologically relevant hemodynamic microenvironments. We observe that like native blood vessels, the local application of force on ECs in our model was dictated by temporal variability in flow and geometric constraints. By defining the pulsatile and oscillatory nature of wall shear stress, our work highlights the dynamic, cyclical nature of shear force experienced by ECs in this model. Furthermore, our novel geometries of stenosis and disturbed flow in this pulsatile, rotational flow system expands the range of hemodynamic environments that can be modeled *in vitro* and provides a scalable platform for investigation of endothelial responses to disturbed flow.

A central goal of this study was to model the long-term adaptation of ECs to complex flow patterns, rather their acute responses to flow onset. Many *in vitro* studies of EC mechanobiology examine responses occurring over a short time frame, ranging from a few hours up to two days. After this period, EC morphology is thought to have stabilized. At the same time, prolonged culture under flow can become challenging since endothelial monolayers in micro- and millifluidic channels can deteriorate. By contrast, we observed stable, healthy, and morphologically stable EC monolayers under rotational flow at five days after initiation of flow. Although proliferation and cell migration were not directly quantified in this study, delaying analysis until five days after flow initiation likely reduced the contribution of these transient processes to the observed phenotypes. Although this duration is far from the years of chronic flow that ECs experience *in vivo*, our work represents an advance toward studying the long-term adaptation of ECs to disturbed flow rather than acute mechanotransduction alone.

CFD provided detailed insight into the complex hemodynamic environments generated within the rotational-flow platform. The simulations revealed that relatively simple geometric modifications fundamentally reshaped local shear environments by altering fluid depth, velocity and flow direction throughout the orbital cycle. These coupled interactions generated local differences not only in WSS magnitude but also in its directionality, pulsatility, oscillatory behavior, and temporal waveform, creating distinct hemodynamic microenvironments that are expected to elicit different endothelial mechanobiological responses. The simulations further demonstrated that relatively simple geometric modifications generated markedly different hemodynamic environments, providing a flexible framework for engineering well-defined hemodynamic microenvironments tailored to specific mechanobiological questions.

The fluid depth used in our experiments was approximately two-fold greater than is typical in EC work. This increased depth was necessary to maintain continuous flow over the obstacle of Case 4. Although greater media depth could theoretically limit oxygen diffusion to the cell layer and lead to EC death,^31^ EC monolayers remained healthy throughout our study. It is likely that in our highly dynamic conditions, vigorous mixing led to enhanced oxygen delivery to the EC layer and minimized diffusion limitations despite increased fluid depth. The maintenance of healthy EC monolayers using these conditions argues that this platform is suitable for prolonged mechanobiology studies.

A notable feature of these simulations was the ability to precisely define the temporal pulsatility of WSS throughout each well. Case 1 and Case 2 both exhibited regions with a rapid dip directly following peak shear, followed by a recovery, producing a waveform that approximates the dicrotic notch observed in the normal aortic pressure waveform. Pulse difference (PD) was most uniform in Case 2, and most heterogeneous in Case 3. The acceleration of fluid generated by the stenosis-like constriction in Case 3, together with downstream flow disturbance, produced highly localized patterns of pulsatility. By contrast, Case 4 exhibited surprisingly uniform PD values, despite substantial variability in the temporal patterns of WSS throughout the rotational cycle.

These observations suggest metrics based solely on WSS levels may fail to capture biologically relevant differences in the temporal WSS exposures. Our findings suggest that regions with the same PD (same min and max values), can nevertheless experience markedly different temporal patterns of shear stress, constituting an additional and largely unexplored mechanobiological variable. Determining whether and how ECs distinguish different WSS patterns throughout the cycle represents an important area for future investigation.

Of note, the pulsatility generated downstream of a stenosis (as in Case 3) is contingent on the compliance of the vessel upstream of the stenosis. A highly compliant upstream compartment would expand to accommodate each pulse of fluid pressure/volume and thereby dampen pulsatility downstream of the stenosis—the so-called “windkessel” effect.^32^ In this scenario, the flow downstream of the stenosis would be *less* pulsatile than the flow upstream. Accordingly, Case 3 may best model the *in vivo* forces present in older, stiffer arteries that do not expand and instead directly transmit the energy of each pulse of flow through the area of stenosis, yielding the dramatic pulse difference seen in our model.

The morphology of ECs is shaped by a multitude of hemodynamic, cellular, and genetic factors. In Case I, the progression of cellular alignment from the outer edge to the center revealed the intriguing finding that ECs align perpendicular to the net WSS vector in regions where both WSS and OSI are high. Prior studies provide limited but important evidence to support this alignment finding. Valvular ECs are exposed to both high shear and high OSI *in vivo*, and exhibit transcriptionally unique patterns, distinct from vascular ECs.^33^ Butcher et al reported the perpendicular alignment of bovine aortic valve ECs to the dominant vector of laminar shear, rather than a simple loss of alignment, and found that this phenomenon disappeared with subsequent passages in culture.^34^ Additionally, one study on impinging flow described perpendicular alignment of ECs to the mean shear vector.^35^ Our data similarly suggest that high WSS/high OSI conditions can activate endothelial transcriptional programs that fundamentally alter the EC mechanobiology. The molecular nature and functional purpose of these adaptations represent an exciting avenue for future investigation.

Abundant literature has characterized the adaptation of ECs to flow *in vitro.* Our study extends existing rotational-flow models by providing detailed spatial maps of WSS magnitude, pulsatility, and oscillatory behavior and correlating these metrics to regional endothelial phenotypes. This low-cost and high-throughput approach, combined and validated with CFD, will empower investigation into the EC responses to the various and complex elements of hemodynamic force *in vivo*.

## MATERIALS AND METHODS

### Details of computational fluid dynamics modeling of rotational flow

To ensure high fidelity of the computational models, detailed structural grids were generated for each model, each containing at least 480,000 elements. As an example, the grid for case 3 contains 837,000 elements with an element size of 0.0005 m. Parameters of all four grids are listed in **Supplementary Fig. 9**.

Computational fluid dynamics (CFD) simulations were performed in ANSYS Fluent, which solves the incompressible Navier–Stokes equations using the finite volume method. A transient pressure-based solver with implicit time integration was employed. Turbulence effects were modeled using the k–ω SST model. Spatial gradients were evaluated using the least-squares cell-based method, while the momentum, turbulent kinetic energy, and specific dissipation rate equations were discretized using second-order upwind schemes.

The air–liquid system was modeled using a volume of fluid (VOF) approach with sharp interface capturing. Two Eulerian phases, air and Dextran-EGM, were included. Surface tension and wall adhesion effects were incorporated to accurately represent free-surface dynamics and fluid–wall interactions. The volume fraction equation was discretized using a compressive scheme to maintain a sharp air–liquid interface.

Orbital motion was prescribed through time-dependent wall velocity boundary conditions to reproduce the motion of the shaker platform. Each geometry was translated in the XY-plane along a circular trajectory corresponding to the orbital motion of the shaker (**Supplemental Video 2**).

Unless otherwise noted, simulations employed a time step of 0.0001 s. The orbital shaker motion was modeled with a radius of 1.9 cm and a rotational frequency of 120 rpm (0.5 s period). Surface tension was modeled using a coefficient of 0.072 N/m, and fluid–wall interactions were represented using a contact angle of 30°. These and other parameters are summarized in **Table 1**.

**Table 1:**
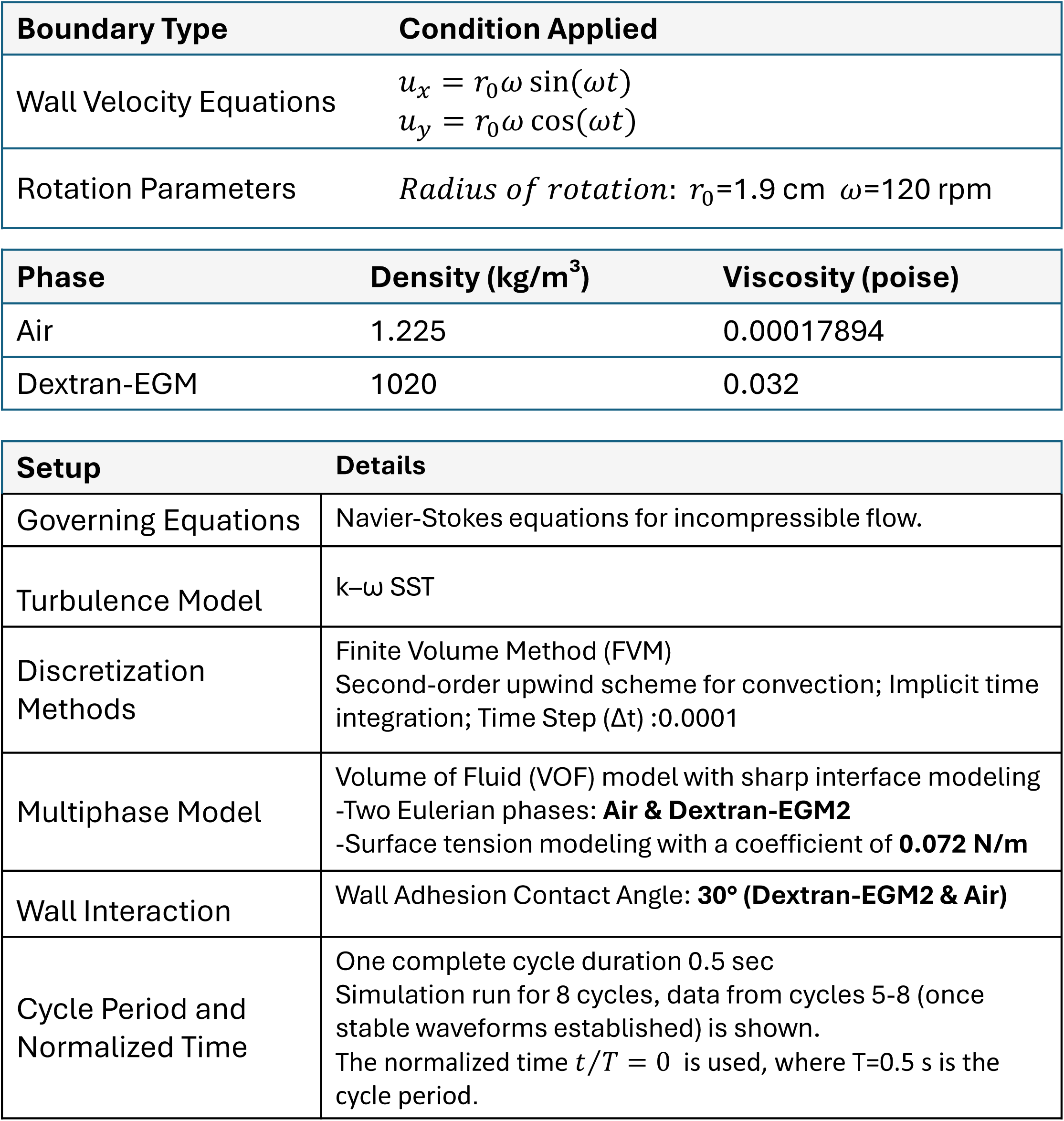
Framework for CFD Analysis.

| Boundary Type | Condition Applied |  |
| --- | --- | --- |
| Wall Velocity Equations | $u_x = r_0 \omega \sin(\omega t)$<br>$u_y = r_0 \omega \cos(\omega t)$ | |
| Rotation Parameters | *Radius of rotation:* $r_0=1.9$ cm $\omega=120$ rpm | |

| Phase | Density (kg/m<sup>3</sup>) | Viscosity (poise) |
| --- | --- | --- |
| Air | 1.225 | 0.00017894 |
| Dextran-EGM | 1020 | 0.032 |

| Setup | Details |
| --- | --- |
| Governing Equations | Navier-Stokes equations for incompressible flow. |
| Turbulence Model | k-ω SST |
| Discretization Methods | Finite Volume Method (FVM)<br>Second-order upwind scheme for convection; Implicit time integration; Time Step ($\Delta t$) :0.0001 |
| Multiphase Model | Volume of Fluid (VOF) model with sharp interface modeling<br>-Two Eulerian phases: **Air & Dextran-EGM2**<br>-Surface tension modeling with a coefficient of **0.072 N/m** |
| Wall Interaction | Wall Adhesion Contact Angle: **30° (Dextran-EGM2 & Air)** |
| Cycle Period and Normalized Time | One complete cycle duration 0.5 sec<br>Simulation run for 8 cycles, data from cycles 5-8 (once stable waveforms established) is shown.<br>The normalized time $t/T = 0$ is used, where T=0.5 s is the cycle period. |

The simulations were initialized with the fluid at rest. To eliminate startup transients as the flow adjusted to the imposed orbital motion, simulations were continued until a repeatable periodic solution was obtained, with negligible cycle-to-cycle variation in free-surface elevation and flow velocity. All reported hydrodynamic quantities were extracted from the final orbital cycle after periodic conditions had been established.

### Magnetic resonance imaging

Representative 2D and 4D flow data from human cardiac MRI was obtained from a single de-identified patient study, processed in Circle (Circle Cardiovascular Imaging, CVI42, v6.1.2). Data was recorded from a 1.5T Phillips MRI scanner (Phillips Ingenia Evolution), as part of the patient’s referral for the procedure under a University of Texas (UT) Southwestern IRB–approved study (STU 032016-009) in 2024.

### Cell culture

Primary human pulmonary artery endothelial cells (HPAECs, Lonza) were grown to confluency in EGM2 media (Lonza) in glass bottom 6-well plates (Cellvis, CA; P06-1.5H-N), modified with plastic inserts. Central inserts were sections of Masterflex® I/P® Precision Pump Tubing (cat no. MFLX96410-26), affixed to the glass bottom of the well with medical adhesive (Factor II, Inc, AZ). The obstacles for models 4 and 5 were longitudinal segments of Nalgene™ Non-Phthalate PVC Tubing (cat no. 8701-4120) cut to 1/8^th^ of the circumference of the tube and similarly affixed to the glass bottom.

For all experiments, cells were rotated at 120 rpm which yielded a cycle time of 0.5 seconds. For rotational flow, we utilized a CO2 resistant shaker (ThermoScientific, cat no. 88881101) inside an incubator maintained at 37°C and 5% CO2.

### Immunofluorescent imaging and quantification

Cells were fixed with 4% paraformaldehyde, washed in PBS, permeabilized in 0.2% Triton/PBS, blocked with CAS-block (Invitrogen) for 30 minutes and then incubated with primary antibody mixture overnight at 4°C, followed by repeat washing and incubation with fluorescent secondary antibodies for 1 hour, followed by DAPI co-staining for nuclear localization. The following were used: anti-CDH5 (, 1:100), anti-GM130 (BD Biosciences, 610823, 1:100), phalloidin-488 (Invitrogen, A12379, 1:400). All secondary antibodies were Invitrogen highly cross-adsorbed secondary antibodies of donkey origin. 6 well plates were imaged on a Nikon CSU-W1 dual-camera inverted spinning-disc confocal microscope, using a custom Jobs module function to scan the entire surface of the well at 10x magnification. A single stitched file was generated using NIS-Elements, and the file converted to .ims format for analysis in Imaris (Oxford Instruments, version 11.0). All experiments were carried out in triplicate and representative images are shown.

For cell alignment analysis, a 2D crop was performed on the region of interest and the Imaris Cells feature as used to identify cells and export data to Microsoft Excel. Cell major axis, cell area, length of cell major axis, and other metrics were calculated by Imaris. CDH5 staining intensity analysis was performed in FIJI, after thresholding the single channel (CDH5) image to isolate the cell membrane. All areas utilized for CDH5 intensity were selected from the middle of an image tile within the stitched image to avoid the mild signal loss at the borders of each field of view. Custom Matlab scripts were used to process and analyze the data generated in Imaris.

### Monocyte Adhesion Assay

To fluorescently label blood monocytes, six million THP-1 cells (ATCC, TIB-202) were incubated in 4 mL of RPMI media containing 5 mL of green Calcein AM (ThermoFisher/Invitrogen, C1430) for 30 minutes, then washed three times with 1x PBS. The fluorescently labeled monocytes were then incubated for 45 minutes with confluent HAECs previously exposed to rotational flow for 5 days. HAECs were subsequently washed to remove non-adherent monocytes. The cells were then imaged using a Nikon CSU-W1 dual-camera inverted spinning-disc confocal microscope. Experiments were carried out in triplicate and the individual data counts from the same region of each plate are shown. Analysis of monocyte adhesion numbers was performed using the Spots feature in Imaris.

### Statistical analysis

Data were both analyzed and plotted in GraphPad Prism 10.2.3. Statistical tests (indicated in the relevant figure legends) utilized were the parametric unpaired Student’s *t* test and the ordinary 1-way ANOVA with Tukey’s multiple-comparison test. Statistical significance was defined as *P* value of less than 0.05. Figures and models were made using Microsoft PowerPoint, data were partially analyzed in Microsoft Excel, and text was written in Microsoft Word.

## ACKNOWLEDGEMENTS

We are grateful to the members of the Cleaver lab for their thoughtful commentary and discussions in the preparation of this manuscript. We appreciate the assistance of the UTSW Quantitative Light Microscopy Core facility in developing the whole-well imaging protocol. We appreciate the assistance of Tarique Hussain, MD/PhD, who provided the representative MRI 4D flow image. The authors acknowledge the University of North Texas High-Performance Computing Services for providing the HPC resources used in this study. This work was supported by the American Thoracic Society/Alveolar Capillary Dysplasia (ACDMPV) Research Grant (23-24PACDA12) to SBS.; the National Institute of Diabetes and Digestive and Kidney Diseases (RC2DK125960, DK106743, DK079862), the National Heart Lung and Blood Institute (HL113498), and the Foundation Leducq grant (21CVD03) to OC.

## AUTHOR CONTRIBUTIONS

SBS, SS, HS, and OC conceptualized the study. SBS designed the cell culture well geometry. SBS, TP, and VSLK carried out cell culture experiments. SBS and MM performed cell imaging and analysis. SS and HS performed all CFD analyses. SBS, SS, HS, and OC provided resources. SBS, SS, HS, OC curated data. SBS and SS generated figures. SBS and OC wrote the original draft of the manuscript, which was reviewed and edited by SBS, SS, VSLK, HS, and OC. SS and OC acquired funding.

## SUPPLEMENTAL FIGURE LEGENDS

**Supplemental Figure 1:**
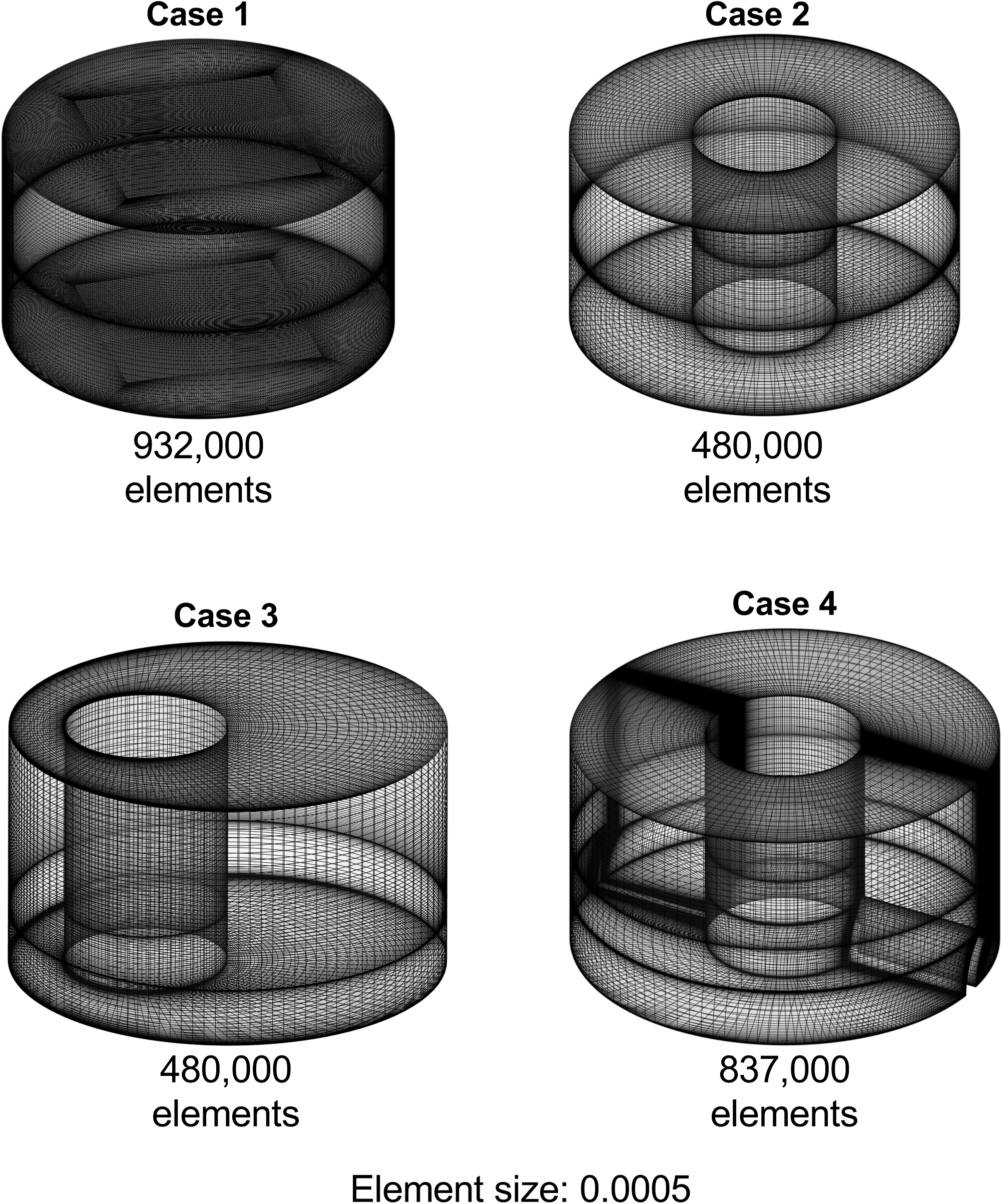
Modeling and grid information for each of the four cases. To improve resolution of oscillatory patterns in Case 1 and 4, a second analysis was done with a higher number of elements. Note that Case 4 was modeled with reverse rotation so that the convex side of the obstacle was upstream and the concave side of the obstacle was downstream.

**Supplemental Figure 2:**
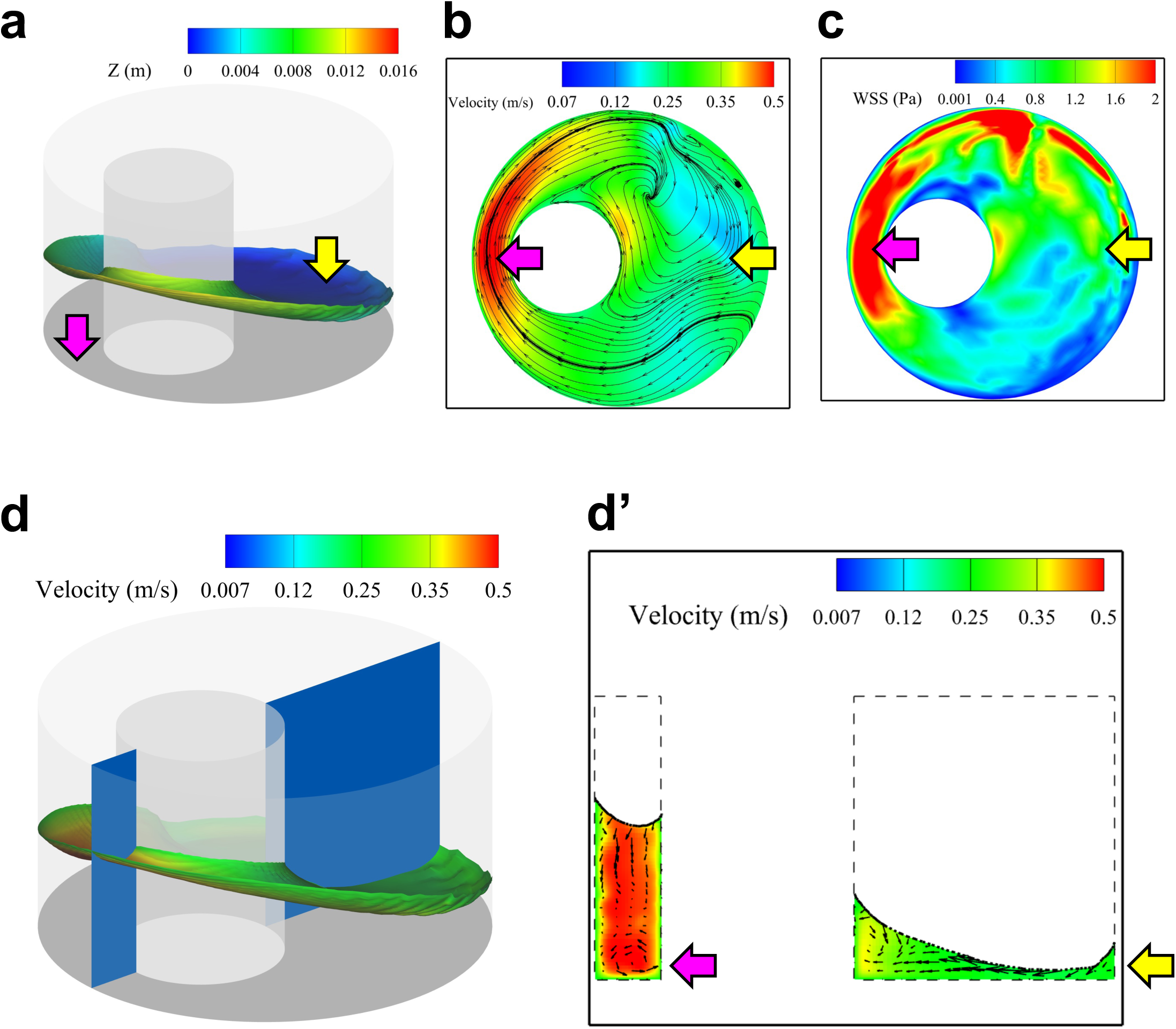
Integration of the effect of fluid depth and velocity in the calculation of wall shear stress. A single time point in the rotational cycle shows the relationship of fluid depth **(a)**, velocity magnitude **(b)**, and wall shear stress **(c)**. The region of flow acceleration through the narrowing gap shows increased velocity (pink arrow), but the reduction of fluid height on the opposite side of the well (yellow arrow) drives shear stress to similar levels, despite the lower velocity. **d,d’)** Cross section of fluid velocity profile at a single point in the cycle reveals vortical flow pattern through the gap with the fluid flow closest to the cell surface directed inward.

**Supplemental Figure 3:**
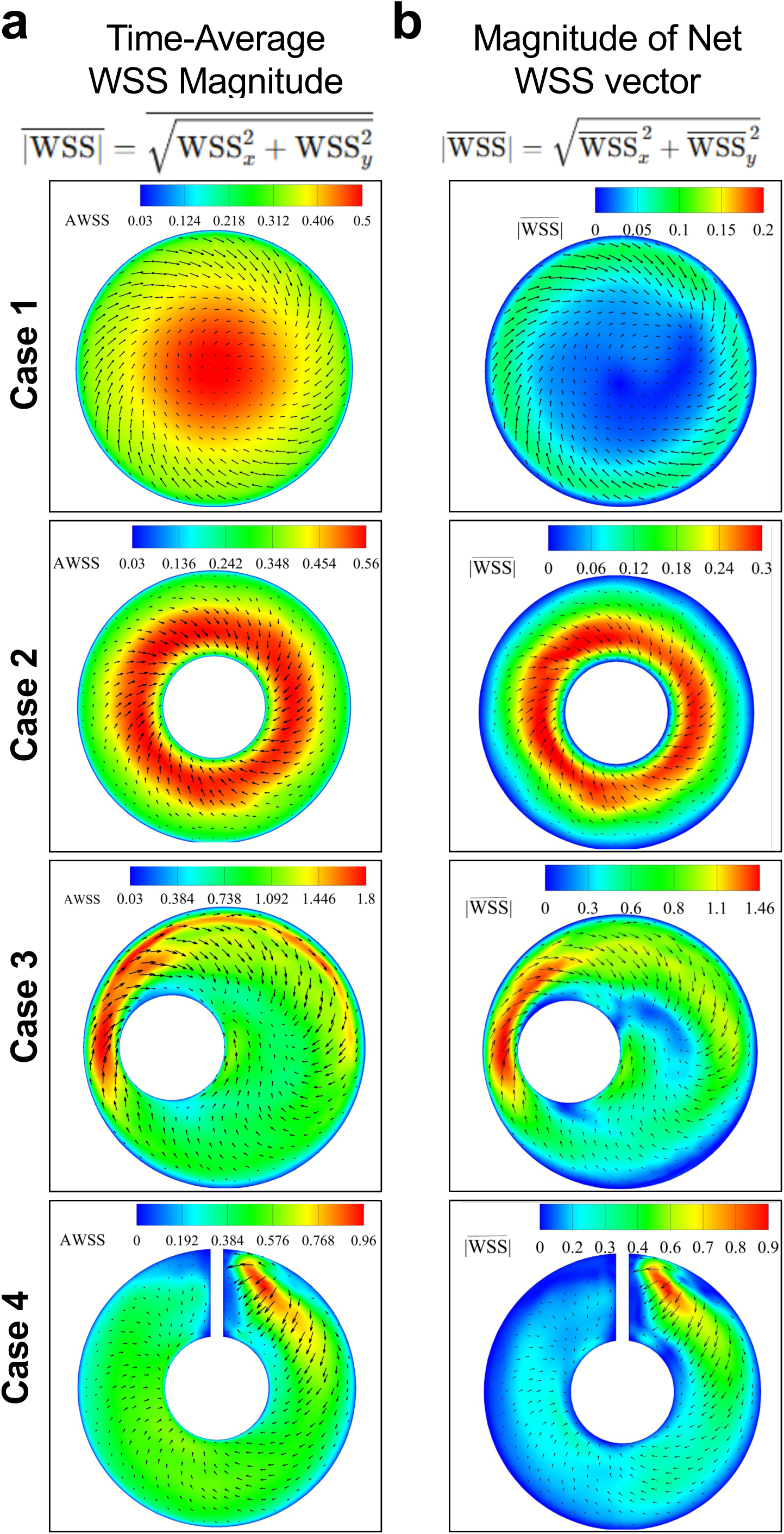
Comparison of average and net WSS values. **a)** Heatmap of average WSS for reach case, with the net WSS vector overlaid. Vector size indicates relative magnitude of the net WSS vector. **b)** Heatmap of the magnitude of the net WSS vector, with the same scaled arrows as overlaid in (a).

**Supplemental Figure 4:**
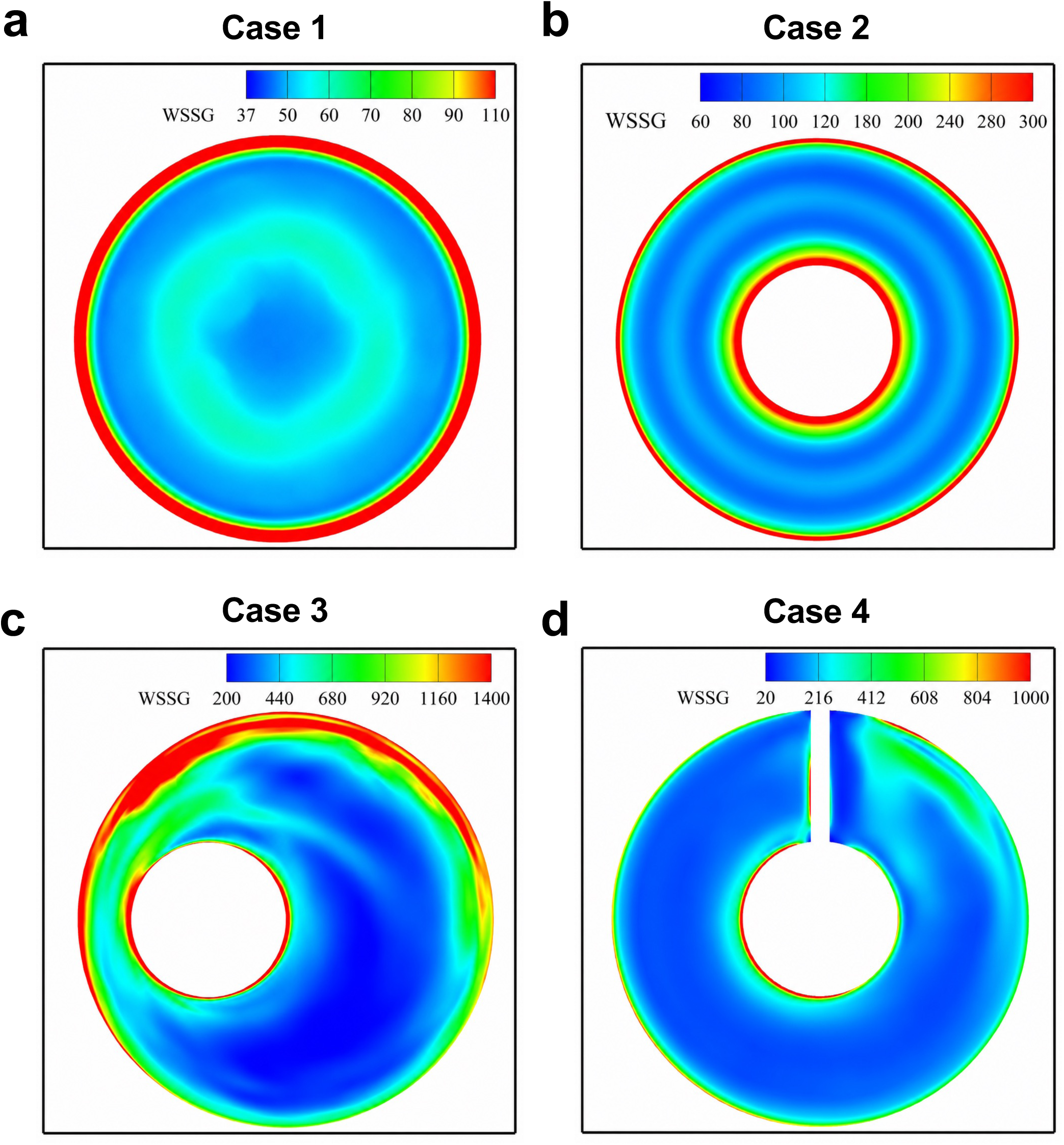
Wall shear stress gradient. **a-d)** Wall shear stress gradient heatmaps for each case.

**Supplemental Figure 5:**
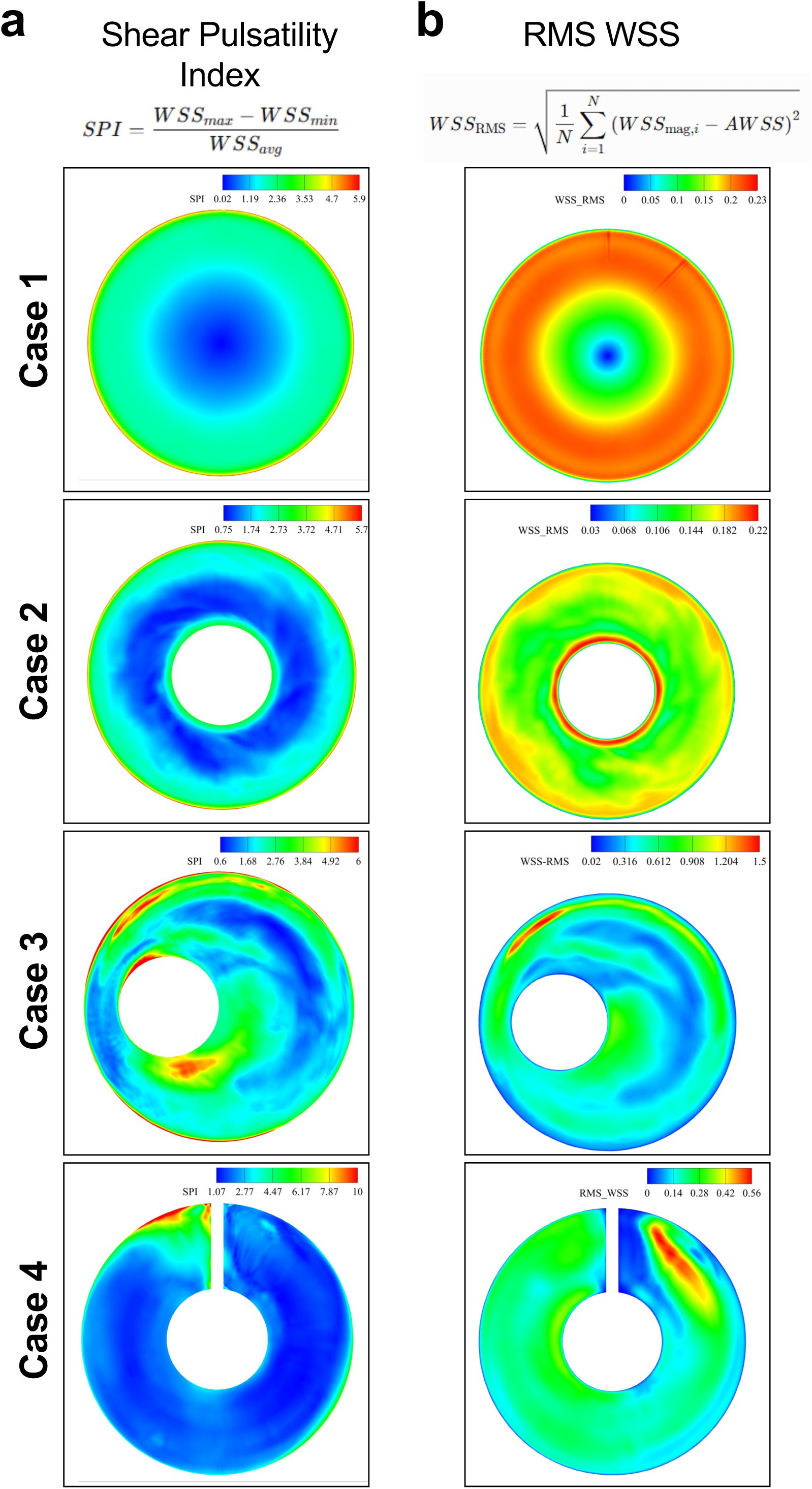
Additional metrics of pulsatility. **a)** Shear pulsatility index for each case. **b)** RMS WSS is another measure of pulsatility that reflects the average magnitude of the WSS oscillations over the entire cardiac cycle.

**Supplementary Figure 6:**
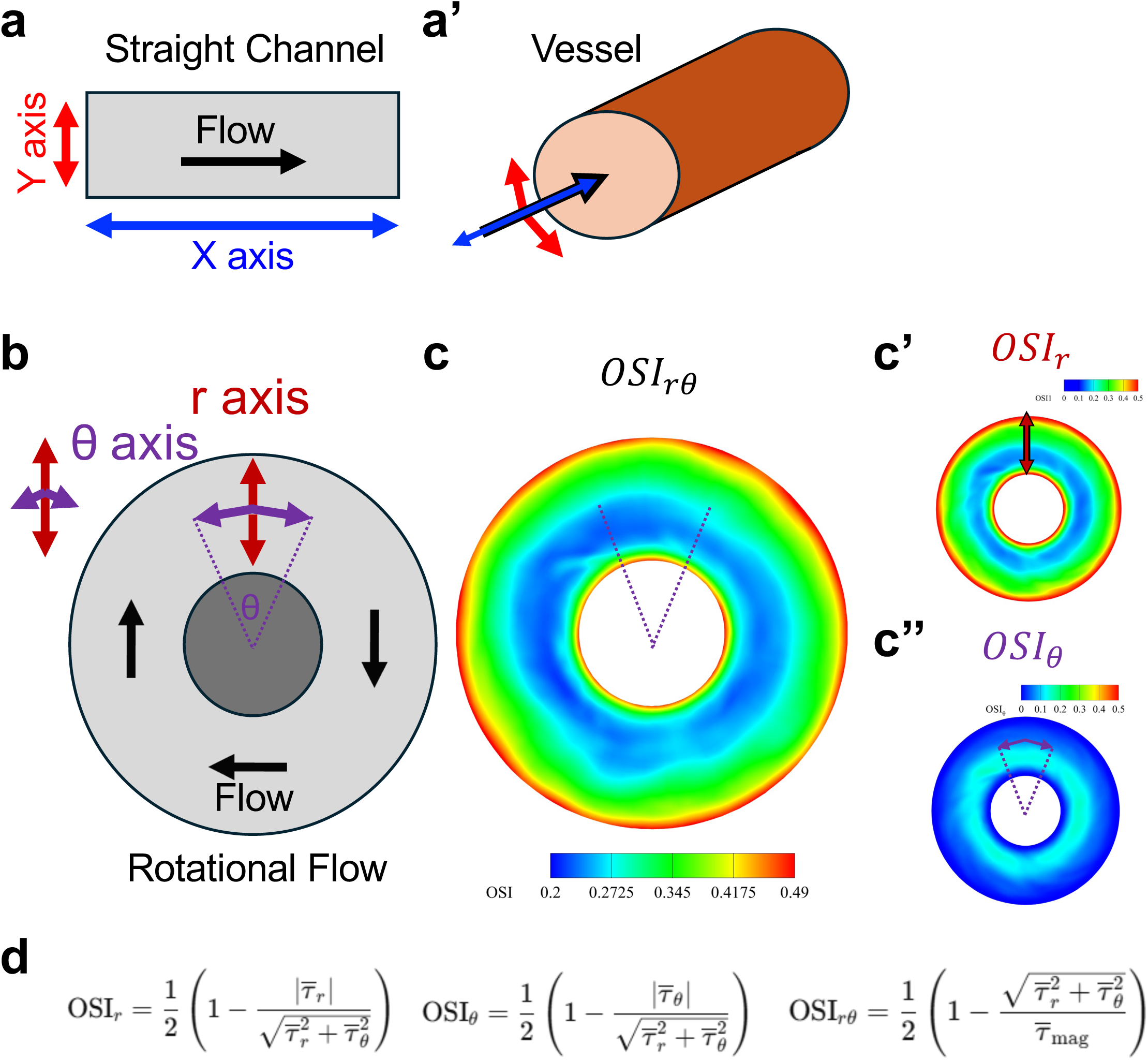
Modeling two axes of oscillatory shear stress in rotational flow assays. **a)** Oscillatory wall shear stress in non-linear vessels on cellular morphology can be considered in two dimensions: along the major axis of the vessel (blue arrows) and in the circumferential axis of the vessel (red arrows). In a simple channel, change in fluid motion/shear stress along can be represented simply along the X and Y axes. In the rotational flow assay, the forward axis is represented by change in radians while the circumferential axis is represented by change in radius. **b, b’)** Computational fluid dynamics modeling heatmaps of oscillatory shear index (OSI) indices in these two dimensions. **(c)** Combined model of OSI incorporating the two dimensions of oscillatory shear. **(d)** Equations governing the color maps in b, b’, and c.

**Supplemental Figure 7:**
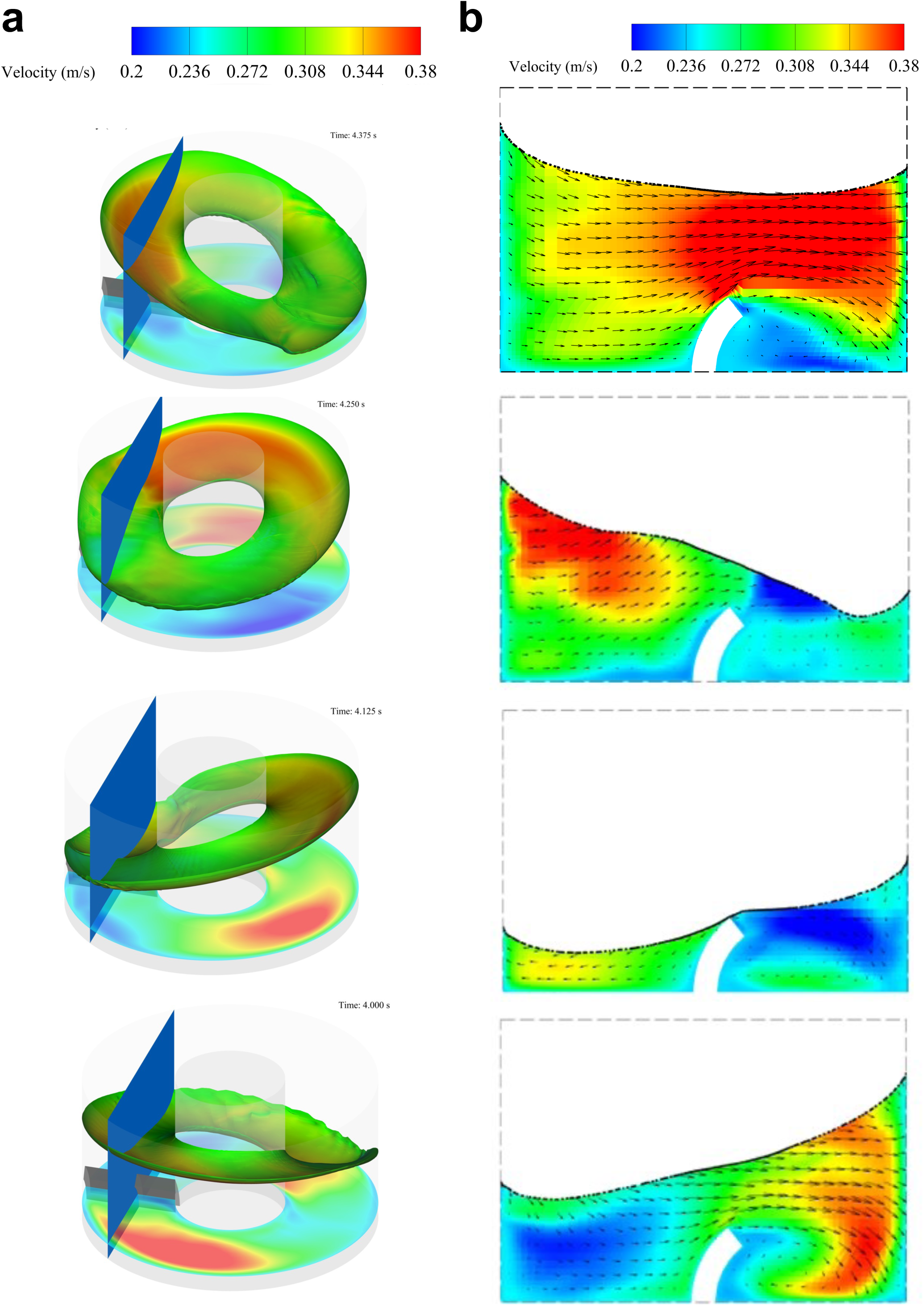
Flow reversal downstream of the obstacle. **a)** Velocity magnitude at air-liquid interface and 2 mm above the surface of the well at four points throughout the rotational cycle. **b)** Velocity magnitude heatmap and streamlines at four points throughout the rotational cycle.

**Supplemental Figure 8:**
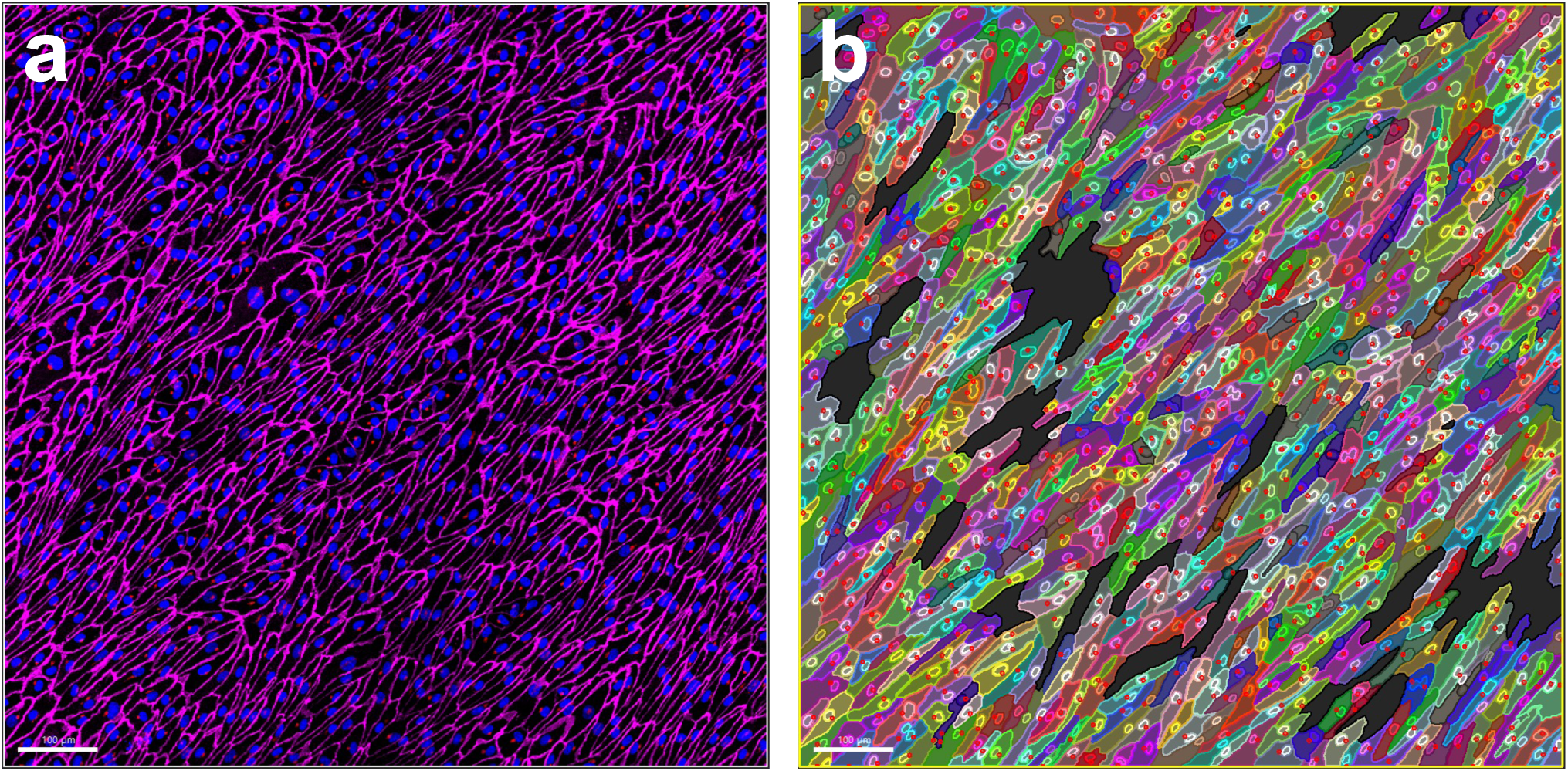
Example mask generated in Imaris. Image shown pre-mask application **(a)** and after application of Cells module mask **(b)**. Gaps represent removal of errors after stringency criteria applied to eliminate as many segmentation mistakes (i.e. multiple cells counted as one) as possible.

**Supplemental Figure 9:**
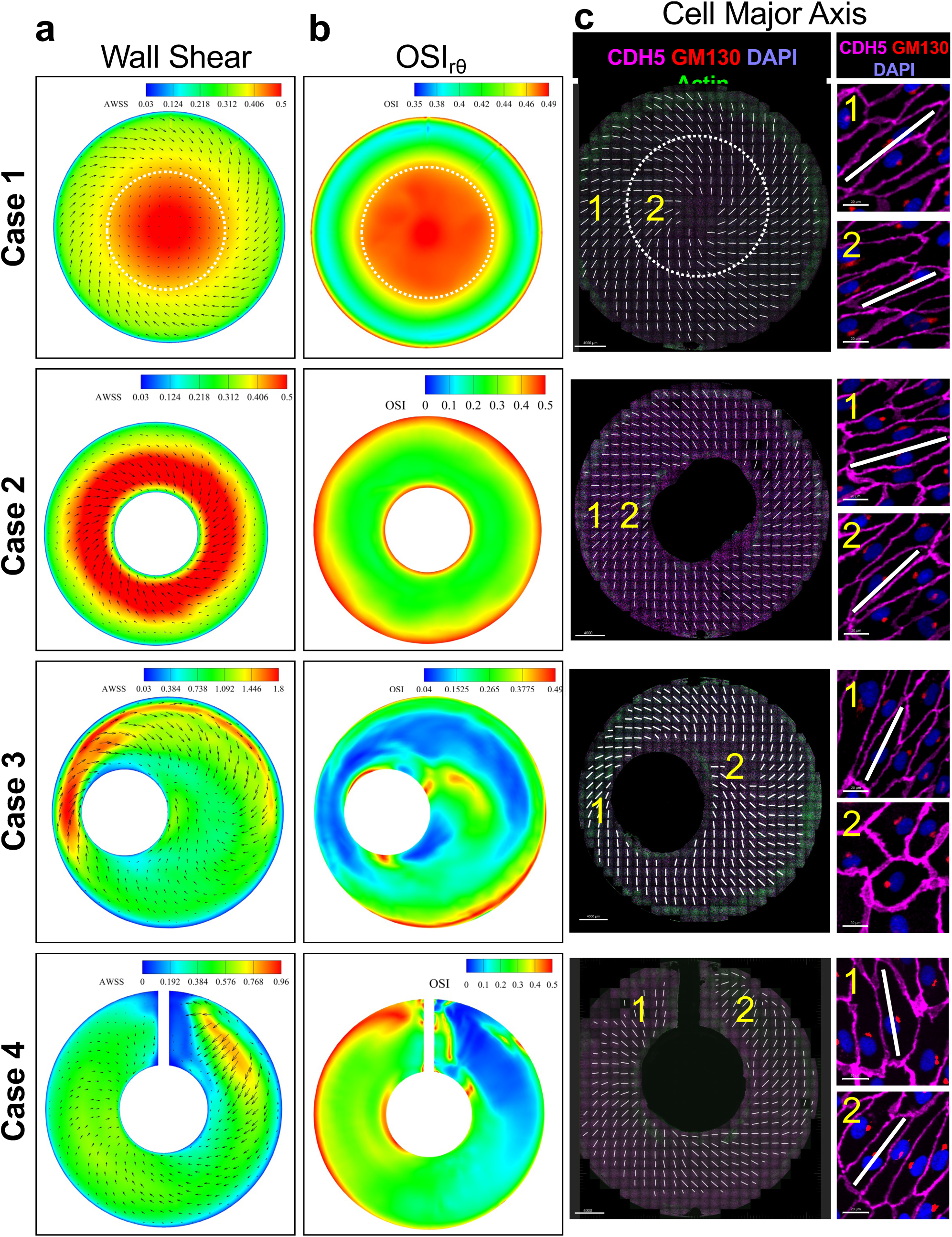
Endothelial cell alignment reflects local WSS direction and oscillatory shear. Juxtaposition of data from Figures 4 and 7 to facilitate comparison and highlight that distinct well configurations (empty, donut, offset, convex obstacle) generate unique patterns of fluid WSS and OSI that influence endothelial alignment. **a)** A heatmap of time-averaged WSS (AWSS) is shown with overlaid vectors indication the direction and magnitude of the net WSS. **b)** Heatmap of time-averaged OSI. **c)** Endothelial cell orientation determined from the major cellular axis following five days of flow exposure. Whole-well scans of IF stains with markers indicated - CDH5 (cell borders), GM130 (Golgi apparatus), and DAPI (nuclei)- with orientation vectors shown as white lines (insets of numbered regions). Example areas (1,2) indicated on low magnification whole well images are shown at high magnification on right, to indicate examples of cells used to determine cell major axes.

**Supplemental Figure 10:**
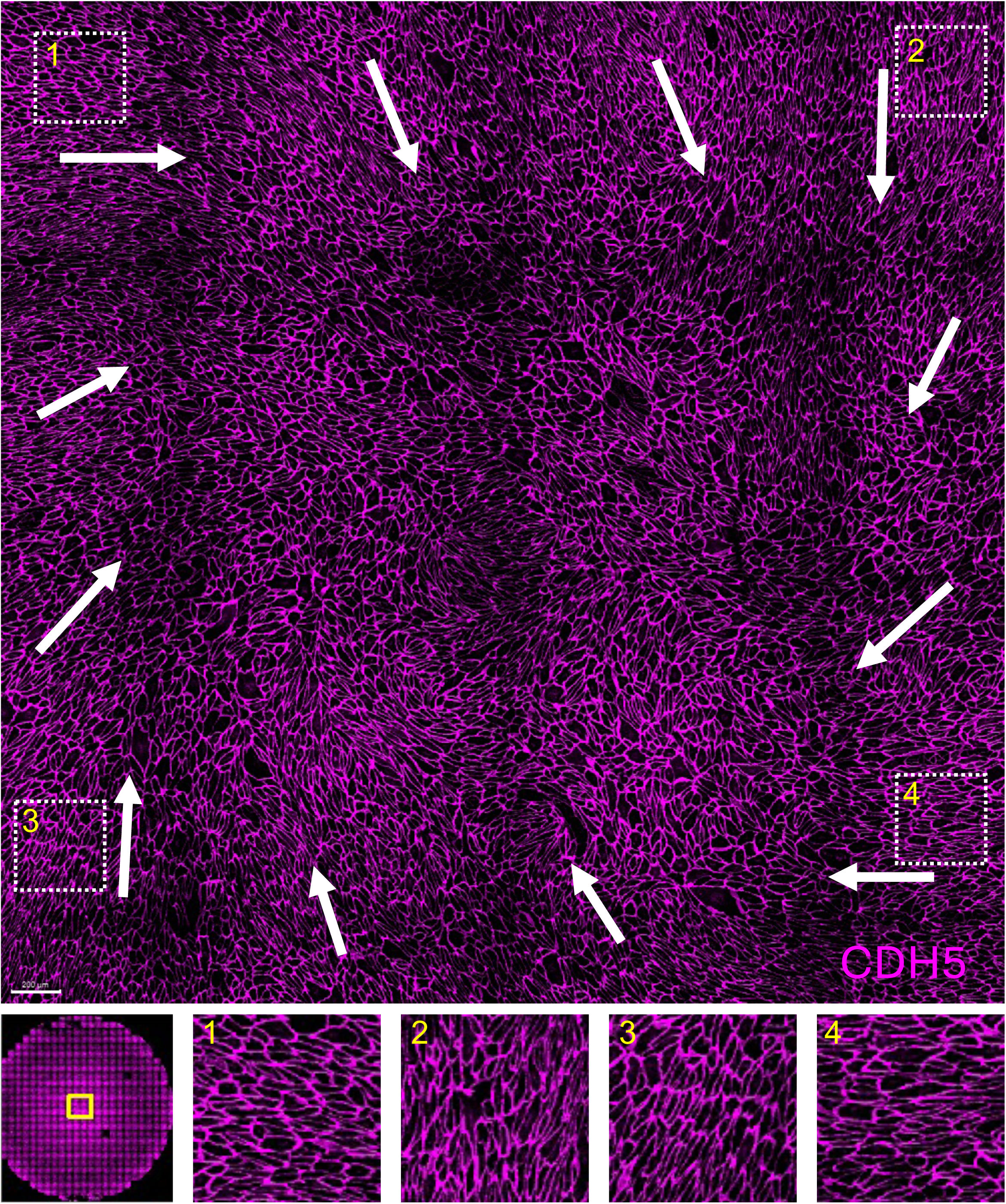
Loss of EC alignment at the center of the open well. Immunofluorescent imaging of cell borders (CDH5) of EC monolayer at the central area of the open well shows the average alignment of the cell major axis (white arrows), with loss of alignment only in the very center of the well. Location within entire well shown by yellow box in lower left. Insets 1-4 show higher magnification of select areas

**Supplemental Figure 11:**
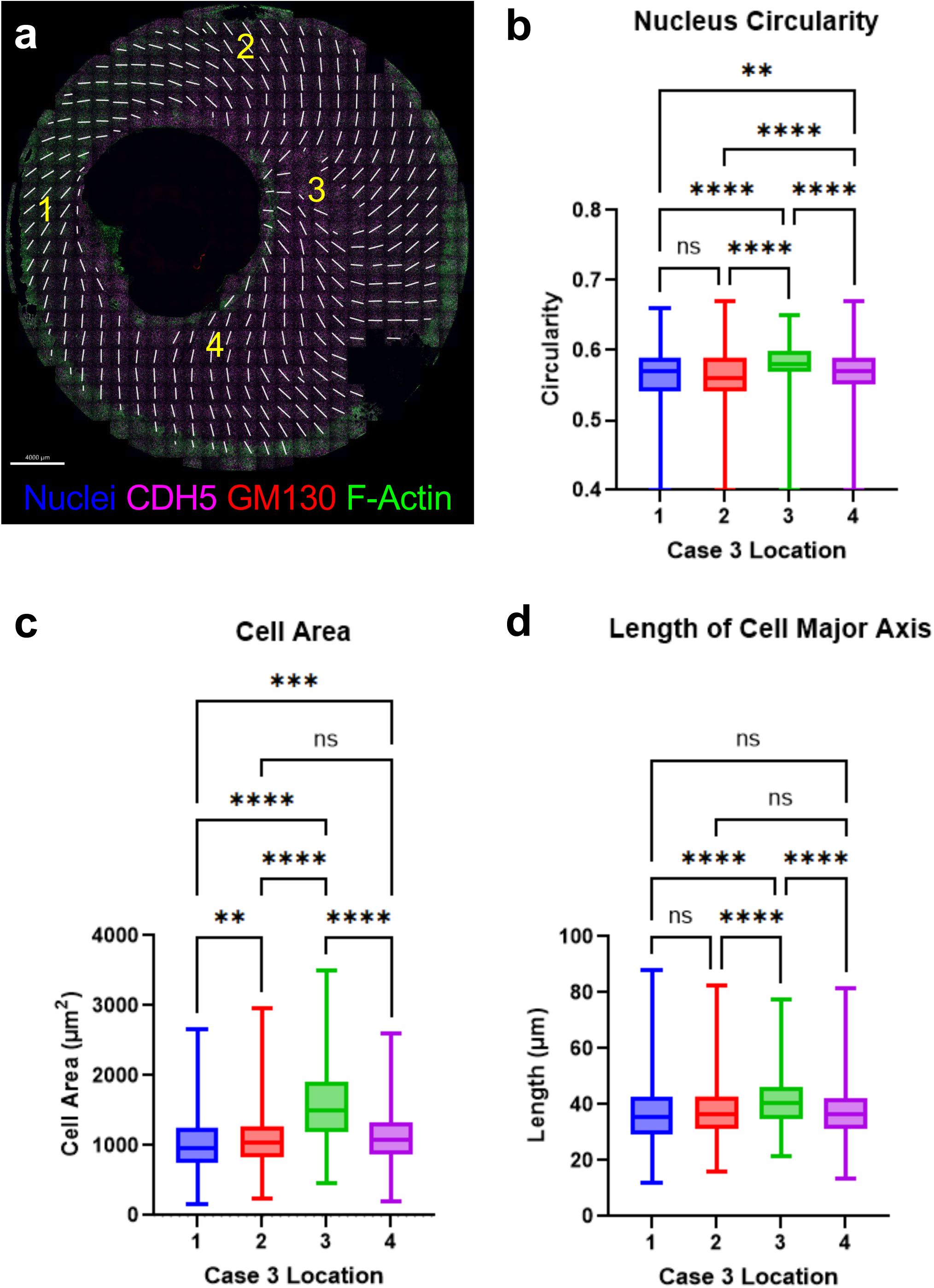
Characterization of cellular and nuclear morphology in the offset well. **a)** selected regions for analysis in the offset well. Quantification of nucleus circularity **(b)**, cell area **(c)**, and length of cell major axis **(d)**.

### SUPPLEMENTAL VIDEOS

**Supplemental Video 1:** Orbital shaker platform with 6-well plate.

**Supplemental Video 2:** Fluid depth changes upon initiation of rotation.

**Supplemental Video 3**: Wall Shear Stress (WSS) changes upon initiation of rotation.

**Supplemental Video 4, 5, 6, 7.** WSS throughout the cycle for each case, after stable waveform is established.

